# Correlated somatosensory input in parvalbumin/pyramidal cells in mouse motor cortex

**DOI:** 10.1101/2022.09.12.507457

**Authors:** Roman U. Goz, Bryan M. Hooks

## Abstract

In mammalian cortex, feedforward excitatory connections recruit feedforward inhibition. This is often carried by parvalbumin (PV+) interneurons, which may densely connect to local pyramidal (Pyr) neurons. Whether this inhibition affects all local excitatory cells indiscriminately or is targeted to specific subnetworks is unknown. Here, we test how feedforward inhibition is recruited by using 2-channel circuit mapping to excite cortical and thalamic inputs to PV+ interneurons and Pyr neurons in female and male mouse motor cortex. Single Pyr and PV+ neurons receive input from both cortex and thalamus. Connected pairs of PV+ interneurons and excitatory Pyr neurons receive correlated cortical and thalamic inputs. While PV+ interneurons are more likely to form local connections to Pyr neurons, Pyr neurons are much more likely to form reciprocal connections with PV+ interneurons that inhibit them. This suggests that Pyr neurons are embedded in local subnetworks. Excitatory inputs to M1 can thus target inhibitory networks in a specific pattern which permits recruitment of feedforward inhibition to specific subnetworks within the cortical column.

**SIGNIFICANCE STATEMENT:** Incoming sensory information to motor cortex (M1) excites neurons to plan and control movements. This input also recruits feedforward inhibition. Whether inhibition indiscriminately suppresses cortical excitation or forms specific subnetworks is unclear. Specific differences in connectivity in circuits promoting different movements might assist in motor control. We show that input to connected pairs of pyramidal (Pyr) excitatory neurons and parvalbumin (PV+) inhibitory interneurons is more strongly correlated than non-connected pairs, suggesting the integration of interneurons into specific cortical subnetworks. Despite sparse connections between these cells, pyramidal neurons are vastly more likely (3x) to excite PV+ cells connected to them. Thus, inhibition integrates into specific circuits in motor cortex, suggesting that separate, specific circuits exist for recruitment of feedforward inhibition.

## INTRODUCTION

Feedforward excitation recruits feedforward inhibition in motor cortex, but whether these inputs silence specific networks faithfully or connect promiscuously is unknown. M1 networks include distinct classes of excitatory pyramidal (Pyr) neurons, organized in layers (Matho et al., 2021; Harris and Shepherd, 2015) as well as inhibitory interneurons. Parvalbumin+ (PV+) interneurons are the largest group of inhibitory interneurons (∼40% of cortical inhibitory cells). PV+ cells are mostly fast-spiking basket cells, targeting the soma and proximal dendrites of excitatory cells, and chandelier cells, though other subtypes exist (Scala et al., 2021; Zhang et al., 2021; Gouwens et al., 2020; Pfeffer et al., 2013; Rudy et al., 2011; Lee et al., 2010; Xu et al., 2010; Kawaguchi and Kondo, 2002; Kawaguchi and Kubota, 1997; reviewed in Tremblay et al., 2016). These excitatory and inhibitory cells receive input from a range of cortical and thalamic sources (Hooks et al., 2013). Somatosensory cortex (S1) projects to topographically defined areas of the corresponding motor cortex (M1) (Aruljothi et al., 2020; Harris et al., 2019; Zingg et al., 2014; Hooks et al., 2013; Izraeli and Porter, 1995; Kaneko et al., 1994). Vibrissal S1 (vS1) most strongly targets L2/3 and L5A neurons in vibrissal M1 (vM1; Mao et al., 2011). Thalamic projections from posterior thalamus (PO), a higher order somatomotor thalamic nucleus, arborize broadly across the tangential surface of cortex. In M1, these axons are laminarly restricted to layers 1 and the border of L2/3 and L5A (Harris et al., 2019; Morgenstern et al., 2016a; Izraeli and Porter, 1995; reviewed in Castro-Alamancos and Connors, 1997), though relative terminal size and density in these layers varies with cortical region (Casas-Torremocha et al., 2019; Audette et al., 2018). Functionally, PO inputs strongly excite L2/3 and L5A Pyr neurons (Hooks et al., 2013; 2015) as well as interneurons (Okoro et al., 2022).

Long-range projections can be cell type-specific (Williams and Holtmaat, 2019; Hu and Agmon, 2016). Both vS1 and PO target Pyr excitatory and inhibitory neurons, with layer-specific complementary activation of PV+ and somatostatin (SOM+) inhibitory neurons in vM1 (Okoro et al., 2022). The organizational principles of cortical neurons studied to date in visual (Palagina et al., 2019; Lee et al., 2016; Morgenstern et al., 2016b; Cossell et al., 2015; Wertz et al., 2015; Glickfeld et al., 2013; Bock et al., 2011; Kätzel et al., 2011; Ko et al., 2014; 2013; 2011; Brown and Hestrin, 2009; Song et al., 2005; Yoshimura and Callaway, 2005; Yoshimura et al., 2005; Gonchar and Burkhalter, 2003; Alonso et al., 2001; Dantzker and Callaway, 2000; Alonso and Martinez, 1998), somatosensory (Naka et al., 2019; Hayashi et al., 2018; Kim et al., 2016; Kätzel et al., 2011; Perin et al., 2011; Brown and Hestrin, 2009; Kampa et al., 2006; Shepherd and Svoboda, 2005; Shepherd et al., 2005; Gibson, J. R. et al., 1999), auditory (Ji et al., 2016; Li et al., 2014; Levy and Reyes, 2012), frontal (Kells et al., 2019; Morishima et al., 2017; 2011; 2006; Kiritani et al., 2012; Kätzel et al., 2011; Hira et al., 2013; Komiyama et al., 2010; Otsuka and Kawaguchi, 2009; 2008; Brown and Hestrin, 2009) and prefrontal (Lee et al., 2014; Wang et al., 2006) cortices suggest existence of subnetworks, a small number of neurons that have higher than random probability of connecting to each other compared to the surrounding cells (Vegué et al., 2017; Buxhoeveden and Casanova, 2002; Mountcastle, 1997). Subnetworks may also share common excitatory inputs or long-range targets (Perin et al., 2013; Yoshimura and Callaway, 2005; Yoshimura et al., 2005; Brown and Hestrin, 2009; Wang et al., 2006). The development of specificity in subnetworks is enhanced by sensory experience in visual cortex (Ko et al., 2014; 2013). This organization may contribute to information propagation and neuronal computation (Peron et al., 2020; Faber et al., 2019; Rost et al., 2018; Nigam et al., 2016).

How inhibitory interneurons integrate in subnetworks may differ from Pyr neurons. Interneurons may connect nonspecifically to Pyr cells in nearby local, intralaminar circuits where they pool inputs from excitatory cells with different response properties (Packer and Yuste, 2011; Fino and Yuste, 2011; reviewed in Fino et al., 2013; Sohya et al., 2007; Mountcastle, Vernon B., 2003).Thus, inhibitory interneurons have broader tuning curves for stimuli orientation and spatial frequency in visual cortex (Bock et al., 2011; Hofer et al., 2011; Niell and Stryker, 2008; Sohya et al., 2007), although inhibitory interneurons show selectivity in some species (Ohki et al., 2005; Hubel and Wiesel, 1963; 1962; but see Ringach et al., 2016; Wilson et al., 2017; Cardin et al., 2007; Hirsch et al., 2003; Moore and Wehr, 2013; Ma et al., 2010; Runyan et al., 2010). How to reconcile specific, connected subnetworks of excitatory cells with the nonspecific targeting of Pyr cells by interneurons? One possibility is that connectivity is dense, with high probability of connection, but synapse strength is weighted higher within subnetworks but strength is weaker to outside networks (Znamenskiy et al., 2018). Another non-mutually exclusive possibility is that long-range inputs and outputs are organized into subnetworks during brain development through Hebbian plasticity (Tezuka et al. 2022; reviewed in Katz and Shatz, 1996).

Here, we examined the functional organization of long-range thalamic (PO) and cortical (vS1) inputs to PV+ and Pyr cells in M1. Here, we refer to these sensory inputs as recruiting feedforward inhibition in M1, though thalamic input to cortex may also be referred to as feedback in other contexts. Using 2-channel optogenetic stimulation with paired whole-cell patch-clamp recording, we show that thalamic and somatosensory inputs were more correlated in connected pairs compared to non-connected pairs. Thus, recruitment of feedforward inhibition by thalamic and somatosensory inputs to motor cortex in mice is subnetwork specific and may depend on functional connections between excitatory and inhibitory cells. Specific differences in connectivity in circuits promoting different movements might assist in motor control.

## MATERIALS AND METHODS

### Animals

Animal protocols were approved by Institutional Animal Care and Use Committee at University of Pittsburgh. Experimental procedures were similar to previous studies (Okoro et al., 2022). Mice of either sex were used at postnatal (P) ages P28-P123 (Average - P46, Median - P43, Mode – P37). PV+-Cre (Jackson Labs, JAX 008069) (Hippenmeyer et al., 2005; Scholl et al., 2015) or SOM-Cre (Jackson Labs, JAX 013044) (Taniguchi et al., 2011) mice were crossed to a lsl-tdTomato reporter line, Ai14 (Jackson Labs, JAX 007914) (Madisen et al., 2010) to label specific interneuron populations.

### Adeno-Associated Virus vectors

AAV2/1.CAG-hChR2-mCherry(H134R).WPRE, titer 1.4E13 (Addgene 100054) (Mao et al., 2011) was injected into posterior thalamus (PO).

AAV2/1.hSyn.ReaChR.mcit.WPRE.SV40, titer 2.52E13 (Addgene 50954) (Lin et al., 2013) was injected into primary vibrissal somatosensory cortex (vS1).

### Stereotactic injections

Animals were anesthetized using isoflurane and placed in a custom stereotactic apparatus. Mice at postnatal day P14-P40 were injected with AAV expressing excitatory opsins. Injections were made with glass pipettes (Drummond) using a custom-made injector (Narashige). The injection apparatus was a positive displacement pump allowing slow injection of nanoliter volumes. Injection coordinates (Table 1) on the anterior/posterior (A-P) axis are reported relative to bregma (positive values anterior to bregma); medial/lateral (M-L) axis coordinates are reported relative to the midline; and dorsal/ventral (D-V) axis coordinates are reported as depth from pia. Injections were made at two depths in cortex. For posterior thalamic injections, we used two adjacent sites, covering the elongated shape (in the A-P axis) of the PO nucleus. The second set of those thalamic anterior injections was done in different mice (n=15) for 11 connected and 12 non-connected PV+ and Pyr pairs. Injections in both sites resulted in similar axon patterns in vM1 (Hooks et al., 2013; reviewed in Castro-Alamancos and Connors, 1997) and were pooled. As in our previous studies, we examined the injection site in thalamus during sectioning to confirm injection targeting to PO. We also confirmed the axonal projection pattern in cortex arborized in layer 1 and the L2/3-5A border, as is typical of PO injections (Hooks et al., 2013; Petreanu et al., 2009).

**Table 1.** Injection coordinates. Anterior/posterior (A-P) axis coordinates are reported relative to bregma (positive values anterior to bregma). Medial/lateral (M-L) axis coordinates are reported relative to the midline. Dorsal/ventral (D-V) axis coordinates are reported as depth from pia. Injections were made at two depths in cortex. Distances in mm.

| Target | A-P | M-L | D-V | Volume |
| --- | --- | --- | --- | --- |
| Posterior thalamus (PO) | -1.3 | 1.2 | 2.5/2.7/2.9 | 50-100nL (x3) |
|  | -0.8 | 1.3 | 2.6/2.75/3.0 | 50-100nL (x3) |
| Primary somatosensory cortex (vS1) | -0.6 | 3.0 | 0.5/0.8 | 50-100nL (x2) |

### Brain Slice Preparation

Brain slices were prepared >14 days after viral injection in young adult mice (P28-P123). Mice were anesthetized with isoflurane and the brain was rapidly removed and placed in cooled oxygenated (95% oxygen and 5% carbon dioxide) choline-based cutting solution (in mM: 110 choline chloride, 3.1 sodium pyruvate, 11.6 sodium ascorbate, 25 NaHCO3, 25 D-glucose, 7 MgCl2, 2.5 KCl, 1.25 NaH2PO4, 0.5 CaCl2). Off-coronal sections (300 µm) of M1 were cut using a vibratome (VT1200S, Leica), rotated slightly from coronal to maintain apical dendrites of Pyr neurons intact in the slice plane. Additional sections were cut to confirm injection location. Slices were incubated at 37 °C in oxygenated ACSF (in mM: 127 NaCl, 25 NaHCO3, 25 D-glucose, 2.5 KCl, 2 CaCl2, 1 MgCl2, 1.25 NaH2PO4) for >30 min and maintained at room temperature (22 °C) thereafter.

### Electrophysiology and photostimulation

Whole cell recordings were performed at 22 °C in oxygenated ACSF with borosilicate pipettes (3-6 MΩ; Warner Instruments) containing potassium gluconate-based internal solution (in mM: 128 potassium gluconate, 4 MgCl2, 10 HEPES, 1 EGTA, 4 Na2ATP, 0.4 Na2GTP, 10 sodium phosphocreatine, 3 sodium L-ascorbate; pH 7.27; 287 mOsm). Data were acquired at 10 kHz using an Axopatch 700B (Molecular Devices) and Ephus software (www.ephus.org) (Suter et al., 2010) on a custom-built laser scanning photostimulation microscope with inversion recovery differential interference (Shepherd et al., 2003) using a Retiga 2000R camera (QICAM; QImaging). Slices were visualized with X4, 0.16 NA, UPlanSApo; Olympus power objective. Individual neurons were visualized with a X60, 1 NA Olympus Fluor LUMPlanFL water-immersion objective. Series resistance errors were minimized with bridge balance in current-clamp mode. Current-clamp recording was performed to confirm stable conditions. To measure excitability, 500 ms current steps were applied starting from -150 pA to 700 pA in 50 pA steps. Connections between pairs of cells were tested in current-clamp with a train of five 3 nA pulses of 0.5 ms duration repeated 40 times (20 for 5 Hz, 20 for 40 Hz) while the other cell was held in voltage-clamp mode. Each sweep length was 2 seconds with a 5 second delay between the sweeps. While testing connections from PV+ interneurons to Pyr cells, the Pyr cell was held at -50 mV (0 mV in some cases – N=5 Pyr cells) to detect IPSCs, while the reciprocal connection was tested with PV+ interneurons held at -70 mV to detect EPSCs.

Photostimulation was done as previously described (Hooks et al., 2015) using 590 and 470 nm LEDs (OptoLED, Cairn). Photostimuli in single-channel experiments were ∼ 2 mW/mm^2^. Photon flux was matched for 590 nm and 470 nm stimuli in the same experiment. Light <585 nm from the 590 nm LED was blocked using a bandpass filter (D607/45, Chroma). Voltage-clamp experiments with LED photostimulation were performed at -70 mV for channelrhodopsin-induced EPSCs and at 0 mV for channelrhodopsin induced IPSCs recruited through feedforward inhibition. Under these conditions, 590 nm LED pulses of 50ms, 100ms, 250ms, and 500 ms were followed by 50 ms pulses of 470 nm LED. Depolarized ReaChR- and ChR2-expressing axons respectively triggered the local release of glutamate. Sweeps were repeated 4 times with a 20 second gap. In some experiments, biocytin was added to the intracellular solution (3 mg/mL biocytin or neurobiotin).

### Histological Preparations and Image Analysis

Some of the slices were processed for biocytin recovery. Samples were postfixed overnight in 4% PFA. Sections were processed as free floating and stained with Hoechst reagent. Blocking was done in TBS containing 10% normal goat serum or normal donkey serum (MilliporeSigma) and 0.5% Triton X-100 (MilliporeSigma) for 2 h at room temperature. Tissue was washed three times in TBS with 2% normal goat serum and 0.4% Triton X-100 (washing solution), followed by incubation with streptavidin conjugated Alexa-647 (1:200) overnight to 24 h at 4°C in the washing solution. After overnight incubation the tissue was stained with nuclear staining Hoecsht (10 mg/mL in water, 1:3,000 dilution; Molecular Probes) for 10 minutes and washed four times for 10 minutes. Tissue was mounted on Fisherbrand ColorFrost Plus microscope slides submerged in Fluoromount G (ThermoFisher).

Images were acquired with a Nikon A1R confocal microscope with X20 or X60 oil-immersion objectives. All images were processed in the ImageJ-Fiji package. Image processing for publication was done in Fiji and Corel Draw Graphics Suite X8 (Corel) or Adobe Illustrator. *Experimental Design and Statistical Analysis.* Data analysis was performed with custom routines written in Matlab. Electrophysiology data were low pass filtered (1 kHz) with an 8-pole low-pass Bessel filter. EPSCs were detected with a threshold of >2x standard deviation from baseline. All data measurements were kept in Microsoft Excel (Microsoft, Redmond, WA) and in Origin (OriginLab, Northampton, MA). Statistical analysis of the data was done in SPSS v.24-v.28 (IBM). For large samples, one-way analysis of variance (ANOVA) with Tukey post hoc correction was used. When the samples had nonhomogeneous variance (significant Levene’s test for equality of variance), Welch’s test with Games–Howell post hoc correction was used. For small samples from different observations, independent-samples two-tailed Student’s t-test was used, and depending on Levene’s test significance, the t statistics for equal or unequal variance are reported. For measurements coming from the same neurons before and after treatment, paired-samples two-tailed Student’s t-test was used. For non-normally distributed data, the non-parametric Wilcoxon signed-rank or Kolmogorov-Smirnov tests were used. All data is shown as arithmetic average ± standard error of the mean (SEM) or ± 95% confidence intervals, unless otherwise specified.

### Code Accessibility

Data analysis was performed with custom routines written in Matlab. Data acquisition and analysis software (M-files in Matlab format) is available upon request.

## RESULTS

To understand how long-range projections to primary vibrissal motor cortex (vM1) excite specific networks of interneurons, we recorded from connected and non-connected pairs of parvalbumin positive inhibitory interneurons (PV+) and Pyr excitatory neurons (Pyr) to explore differences in their circuit connectivity. We used the 2-channel ChannelRhodopsin-Assisted Circuit Mapping (2CRACM) approach developed by our lab (Hooks et al., 2015; Petreanu et al., 2009; Petreanu et al., 2007). We injected viral vectors containing Channelrhodopsin-2 (ChR2-mCherry) into posterior thalamus (PO) and the red-shifted ChannelRhodopsin variant ReaChR (ReaChR-mCitrine) into vS1. We activated these opsins by sequential 590 nm and 470 nm stimulation (Fig. 1A,B,J,K). The intracranial injections were done with a custom-made positive displacement system in PV+-Cre^+/+^;lsl-tdTomato (ai14)^+/+^ mice. To allow for opsin expression, recording started two weeks after injections. Pairs of adjacent (< 120 µm) PV+ and Pyr neurons were recorded in whole-cell current- and voltage-clamp configuration. Passive membrane properties and series resistance were measured at -70 mV. Current injections of 500 ms in 50 pA steps characterized active membrane properties including Action Potential (APs) firing (Fig. 1E,H; Supp. Fig. 1; Supp. Table 1). Connectivity was tested in each direction (PV+ ↔ Pyr), holding one neuron in current-clamp and applying 3 nA, 0.5 ms current steps while the other neuron was held in voltage-clamp (Fig. 1F, I). Some of the slices were processed for biocytin recovery at the end of the experiments, which allowed confirmation of cell type and laminar position (Fig. 1D,G). Input was quantified in voltage-clamp using the 2CRACM approach, ReaChR-mCitrine was stimulated with 590 nm LED light (50-500 ms pulses), followed by stimulation of ChR2-mCherry with 470 nm blue LED light (50 ms pulses immediately following), with additional 500 ms 590 nm only LED stimulation to have ReaChR trace only for subtraction (Fig. 1J,K).

**Figure 1.**
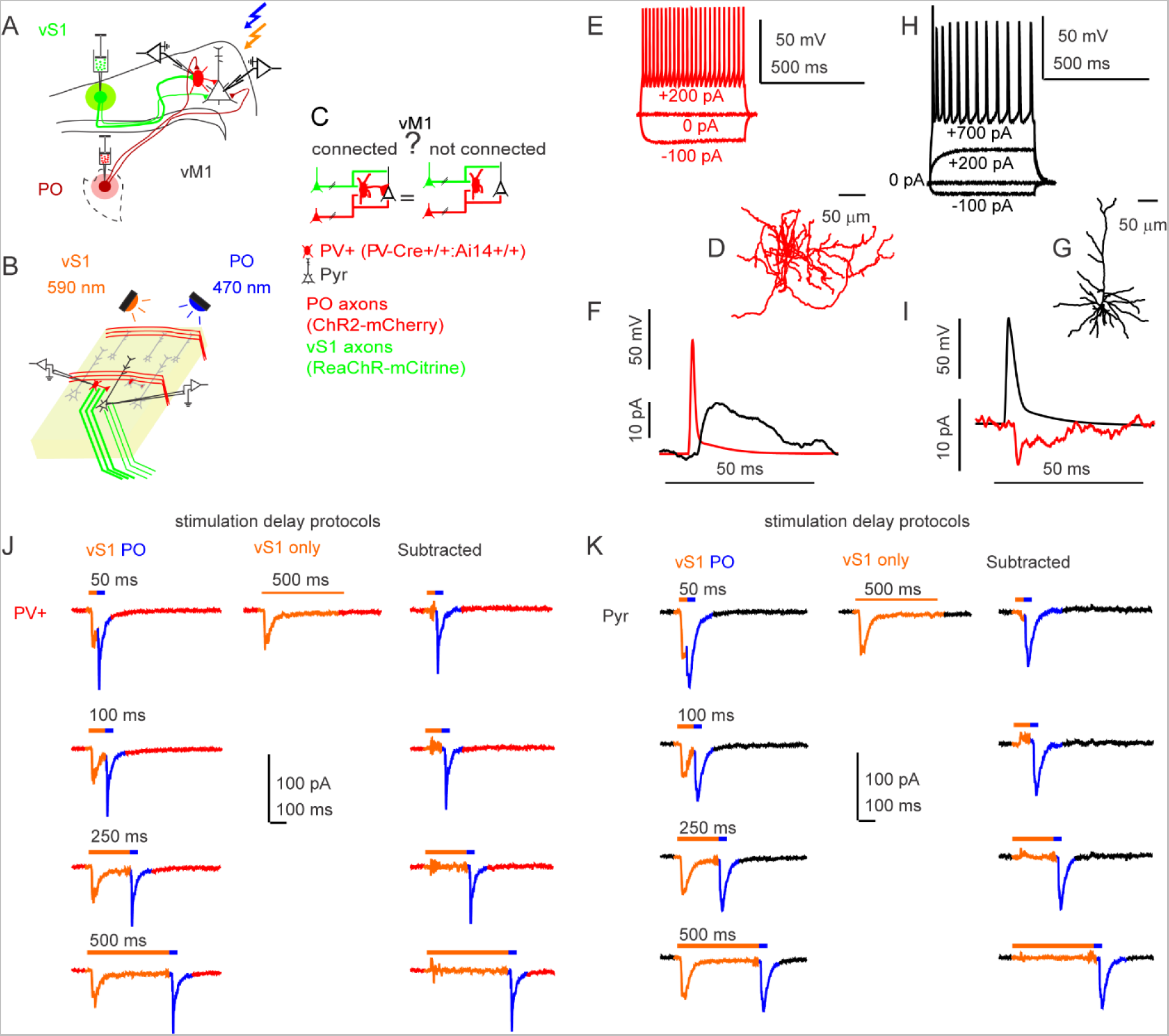
Long-range thalamic and cortical projections in mouse brain slices. **A.** Illustration of the slice preparation. Vibrissal M1 (vM1) receiving long-range protection inputs from vS1 (green lines, ReaChR-mcitrine expressing), and from posterior thalamus (red lines, ChR2-mcherry expressing). Paired whole-cell patch-clamp recording targeted to a pyramidal excitatory neuron (Pyr, empty triangular shaped) and parvalbumin positive inhibitory interneuron (PV+, oval shaped, red) receiving inputs from vS1 (stimulated with orange LED, 590 nm), and from posterior thalamus (stimulated with blue LED, 470 nm). **B.** Illustration of stimulation paradigm in brain slices. ReaChR expressing axons (vS1, green) were stimulated first with 590 nm, orange LED (50-500 ms) immediately followed by stimulation of ChR2 containing axons (PO) with 470 nm blue LED 50 ms with equal light intensity (∼2mW/mm^2^). Example traces are shown in (J-K). **C.** Illustration of scientific inquiry question and color coding. **D.** Example reconstruction of a recorded biocytin filled PV+ cell. Scale bar above. **E.** Example current-clamp traces showing responses of a PV+ inhibitory fast spiking cell recorded in those experiments with current steps in between and the scale bar to the right. **F.** Example traces of a connectivity test between a PV+ cell in current-clamp mode with 3 nA 0.5 ms current step to elicit single AP and the voltage-clamp IPSC response of the Pyr cell held at 0 mV. **G.** Example reconstructed biocytin filled Pyr cell. Scale bar above. **H.** Example current-clamp recording of Pyr cell with current steps as labeled. Scale bar to the right. **I.** Example of connectivity test, with the same protocol as in (E), Pyr in current-clamp, EPSC in PV+ cell held at -70 mV. **J-K.** Examples of two connected PV+ and Pyr cells in voltage-clamp showing responses to LED stimulation. First column 590 nm and 470 nm LED stimulation (colored bars indicate time of LED on/off). Middle traces are 500 ms 590 nm alone. Third column is a subtraction of middle traces from the first column to reveal 470 nm response.

The channelrhodopsin-induced EPSCs kinetic properties are shown in Fig. 2. These include the EPSC onset delay, the rise time, normalized amplitude, and decay time course. EPSC onset times may be slightly slower for vS1 inputs compared to PO inputs, due to the slower kinetics of ReaChR (Lin et al., 2013). However, these do not vary significantly across delay times, suggesting that activating vS1 inputs first does not affect responses from PO afferents.

**Figure 2.**
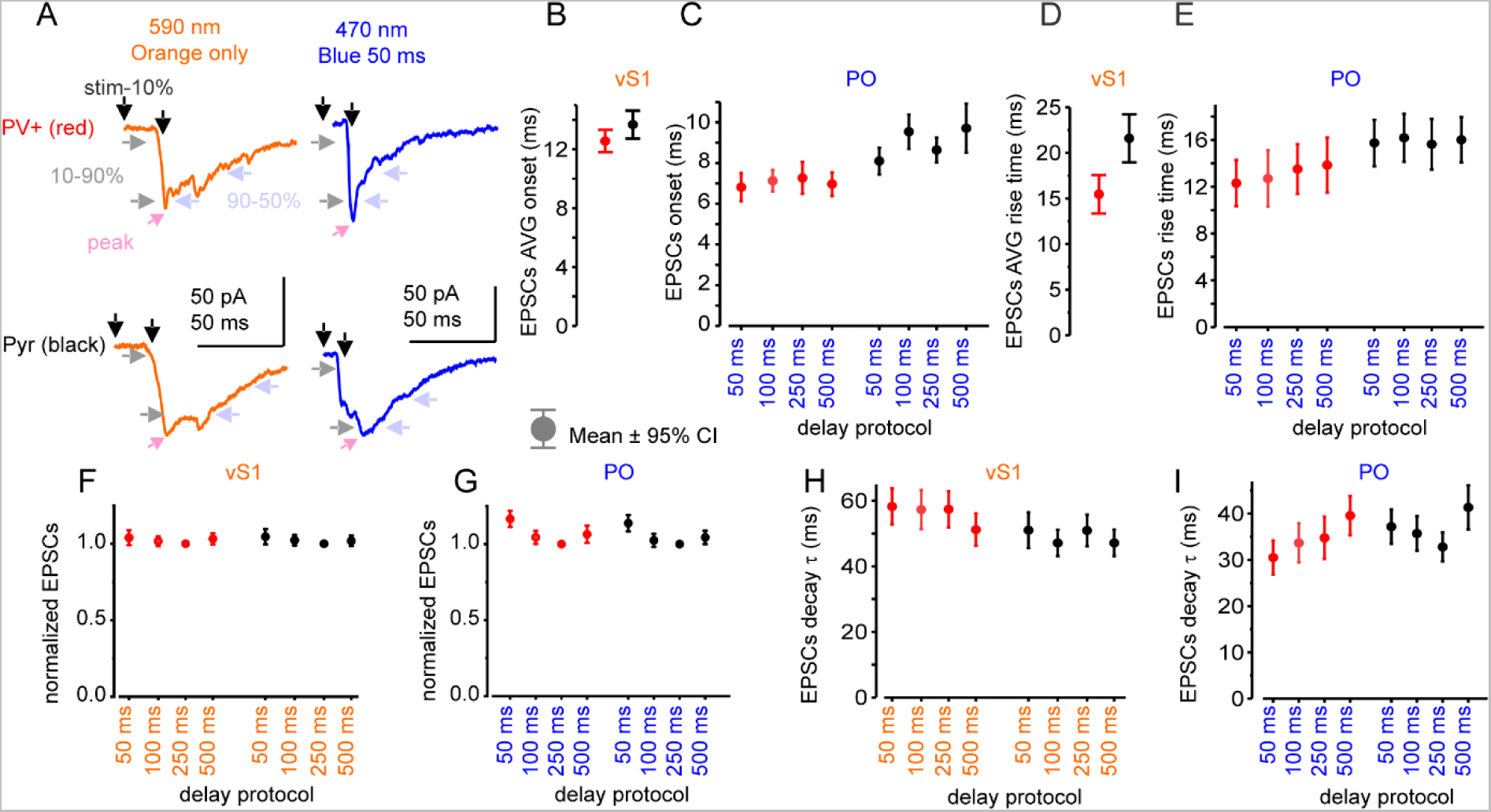
2CRACM EPSCs kinetics. **A.** Example response traces of PV+ (red) and Pyr (black) to 590 nm (vS1, orange) 500 ms LED stimulation, following by 470 nm (PO, blue) 50 ms LED stimulation. Description of kinetics that were measured and compared, as indicated by arrows in the panel. Onset was measured from start of the stimulation to 10% of the EPSC peak, the rise time was measured from 10% to 90% of the EPSC peak; and the decay time was measured from 90% to 50% of the EPSC peak. Peaks are shown by pink arrowheads. Because vS1 responses onset prior to PO responses, onset and rise kinetics are averaged across all 4 delay protocols. **B.** vS1 EPSC onset averaged across all delay protocols (50-500 ms), (PV+ n=99; Pyr n=99). **C.** PO EPSC onset (PV+ n=52-66; Pyr n=59-83). **D.** vS1 EPSC averaged rise time (PV+ n=103; Pyr n=103). **E.** PO EPSC rise time (PV+ n=62-80; Pyr n=72-95). **F.** vS1 EPSC normalized to its own peak at 250 ms delay protocol response (PV+ n=101-104; Pyr n=111). **G.** PO EPSC normalized to its own peak at 250 ms delay protocol response (PV+ n=71-87; Pyr n=87-104). **H.** vS1 EPSC decay time (PV+ n=37-82; Pyr n=42-86). **I.** PO EPSC decay time (PV+ n=52-69; Pyr n=67-85). Means are shown with 95% confidence intervals.

### Somatosensory cortical excitation is stronger than thalamic input to PV+ neurons in layer 2/3 of vM1

To compare the difference in recruitment of feedforward inhibition mediated by PV+ cells and excitation of Pyr cells, we recorded opsin-mediated EPSCs in pairs of PV+ and Pyr cells in the whole-cell voltage-clamp configuration. Cortical laminae were defined based on visible boundaries formed by differential cell densities in the brightfield image of the slice (Weiler et al., 2008; Hooks et al., 2011) and reported as the normalized distance of the cells between pia and white matter. L5A is the pale band in the brightfield image above the more heavily myelinated L5B (Yu et al., 2008). L2/3 cells were within 8-38% and L5A cells were within 20-58% of the slice thickness, depending on the curvature and anterior-posterior position of the slice (Fig. 3C). Our prior data, using subcellular channelrhodopsin-assisted circuit mapping (sCRACM), had shown that both vS1 input and PO input similarly excited vM1 L5A Pyr neurons most strongly and L2/3 Pyr neurons with ∼70-90% of this strength (Mao et al., 2011; Hooks et al, 2013). But for PV+ neurons, this pattern shifted and vS1 excited L2/3 PV+ neurons more strongly than L5A, while PO excited L5A PV+ neurons more strongly than L2/3 (Okoro et al., 2022). Based on this difference in connection strength, EPSCs amplitudes in PV+ cells divided by the amplitudes of EPSCs in Pyr cells should result in larger ratios from vS1 stimulation compared to PO stimulation in L2/3 in our paired recordings. Consistent results confirm that the wide-field LED stimulation in the 2CRACM approach produces similar results to those predicted from earlier circuit mapping approaches (which differs by the use of tetrodotoxin (TTX) to prevent action potentials in the slice and ensure monosynaptic responses; Petreanu et al., 2009). We grouped recordings by laminar position, focusing on pairs within L2/3 and L5A, where long-range input from the pathways studied is strongest.

**Figure 3.**
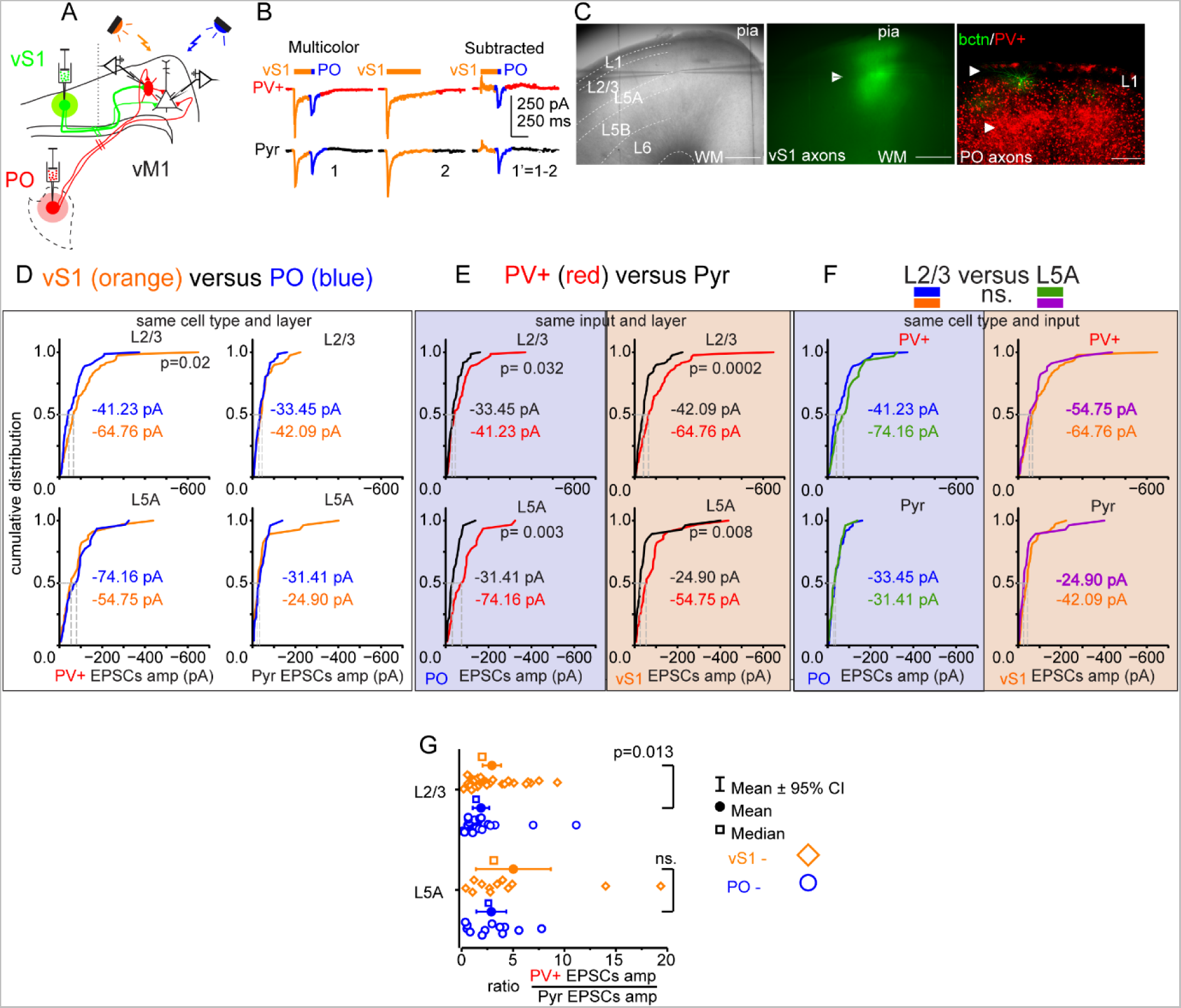
vS1 input is larger than PO input to vM1 layer 2/3 PV+ interneurons. **A.** Illustration of the vM1 slice preparation receiving long-range projections inputs from vS1 (green lines, ReaChR-mcitrine) and from PO (red lines, ChR2-mcherry) during paired recording. **B.** Example voltage-clamp traces in cells held at -70 mV from a pair of PV+ inhibitory interneuron and Pyr excitatory neuron showing responses to 590 nm LED stimulation of vS1 axons in vM1 immediately followed by 470 nm LED stimulation of PO axons. Middle panel is the response to 590 nm LED stimulation alone and the last panel is a subtraction of the middle traces from the first traces to isolate 470 nm-induced EPSCs. **C.** Example of an off-coronal 300 µm brain slice of vM1 with two cells in patch-clamp. Left panel shows the location of the cells relative to pia and white matter. Approximate layer boundaries indicated. Top right panel shows vS1 axonal fluorescence in green (arrowhead). Lower panel is a confocal image showing PO axonal fluorescence in red (arrowheads) in L1 and the lower L2/3 and L5A together with PV+ interneurons. A biocytin filled example pair (green) of cells is shown. Scale bars are 500 µm. **D.** Cumulative distribution of EPSC amplitudes comparing vS1 and PO inputs to PV+ (left) and Pyr (right) neurons in layers 2/3 (top) and 5A (bottom). EPSCs in L2/3 PV+ neurons were significantly larger from vS1 than PO (Mann-Whitney U=3596, p=1.97E-2). Differences between vS1 and PO EPSCs in L2/3 Pyr neurons were not significant (Mann-Whitney U=3009, p=0.44). PV+ and Pyr EPSCs amplitudes in L5A were not significantly different (PV+: Mann-Whitney U=468, p=0.44; Pyr: Mann-Whitney U=334, p=0.6). **E.** Cumulative distribution of EPSC amplitudes comparing input to Pyr and PV+ neurons from PO (left) and vS1 (right) to neurons in layers 2/3 (top, M-W U=2151 and 2025 (PO), p=2.34E-4 and 3.2E-2 (PO)) and 5A (bottom, M-W U=289 and 217 (PO), p=8.17E-3 and 2.88E-3 (PO)). **F.** Cumulative distribution of EPSC amplitudes comparing L2/3 and L5A inputs to PV+ (top) and Pyr (bottom) from PO (left) and vS1 (right) was not significantly different (PO L2/3 to L5A PV+: M-W U=1369, p=0.069; L2/3 to L5A Pyr: M-W U=897, p=0.83; vS1 L2/3 to L5A PV+: M-W U=1257, p=0.41; L2/3 to L5A Pyr: M-W U=883, p=0.11). **G.** Ratio of vS1 and PO EPSCs amplitudes in PV+ interneurons divided by the EPSCs amplitudes in Pyr neurons (from D-F) shows that vS1 preferentially targets PV+ neurons compared to PO confirming the results of previous study with subcellular CRACM (Okoro et al., 2022). Only pairs with both inputs included. L2/3 vS1 and PO pairs, n=30; L5A, n=12. Wilcoxon signed-rank test (Wlcxn srt =2.48, p=1.3E-2) was used since the synaptic responses are not normally distributed Kolmogorov-Smirnov and Shapiro-Wilk tests for normality (L2/3 PO inputs (PV+/Pyr) K-S(12)=0.312, p=2.02E-3, S-W(12)=0.7, p=8.35E-4; L5A vS1 inputs (PV+/Pyr), K-S(12)=0.34, p=4.15E-4, S-W(12)=0.718, p=1.26E-3. Means are shown by circles and the medians by squares. Vibrissals represent 95% confidence intervals.

EPSC amplitudes showed that vS1 inputs to PV+ cells are stronger compared to PO inputs in vM1 L2/3 (Fig. 3D, Mann-Whitney, p=1.97E-2 for EPSCs), while comparisons in L5A showed similar synaptic strength (Fig. 3F,G). Comparisons of input strength to PV+ and Pyr neurons in the same layer generally revealed stronger amplitude EPSCs to PV+ neurons (Independent-Samples Mann-Whitney test, vS1 inputs to L2/3 PV+ compared to Pyr, total N=161, p= 2.34E-4; to L5A PV+ compared to Pyr, total N=62, p= 8.17E-3; PO inputs to L2/3 PV+ compared to Pyr, total N=143, p= 3.2E-2; to L5A PV+ compared to Pyr, total N=57, p= 2.88E-3.) As predicted, the PV+ to Pyr input ratio (Fig. 3G) was greater for vS1 inputs than for PO inputs (Wilcoxon signed-rank, p=1.3E-2).

### Connectivity between the PV+ and Pyr neuron pairs

Paired recordings allowed identification of Pyr and PV+ neuron pairs that were connected or unconnected, providing comparison between neurons in the same local network. Connectivity was tested bidirectionally between 197 PV+ and Pyr pairs (Fig. 4A-C). Of these, ∼39% were connected (N=77/197), either unidirectionally or bidirectionally.

**Figure 4.**
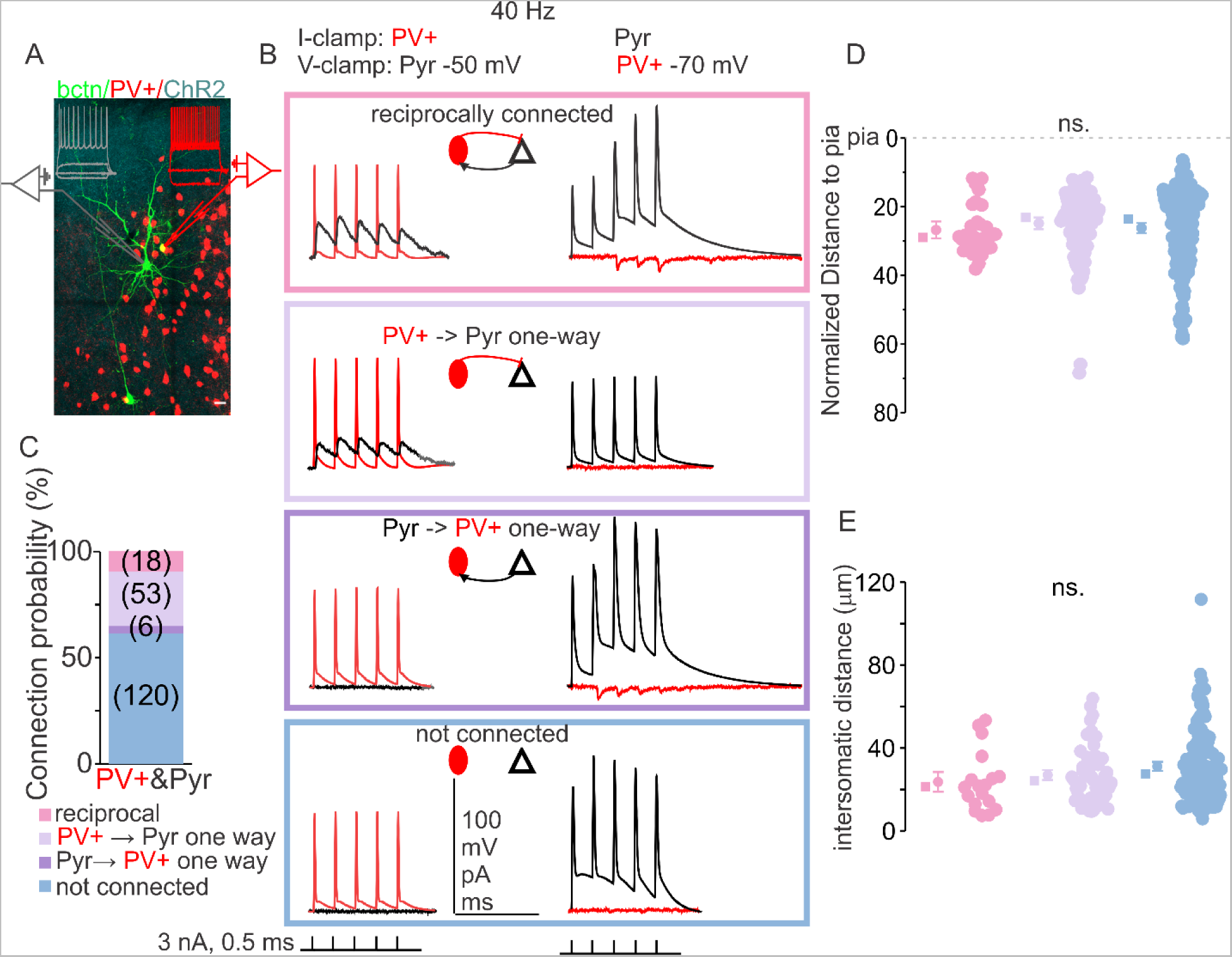
Characterization of connectivity between PV+ and Pyr neurons. **A.** Example of biocytin filled 3 cells (green) and current-clamp traces show characteristic PV+ (red) and Pyr neurons subthreshold responses and suprathreshold AP firing in response to depolarizing current steps (PV+ red, Pyr gray) with ChR2 positive axons (cyan). One PV+ filled with biocytin is shown in yellow, due to overlap of tdtomato labeling. **B.** Example of connection test traces for reciprocally connected (top), connected in one-way from PV+ to Pyr (2^nd^ row), connected in one-way from Pyr to PV+ (3^rd^ row), and non-connected (4^th^ row). **C.** Percentage of connected pairs (Fisher’s Exact 20.81, p=1.6E-5). **D.** Normalized distance to pia, the distance from pia to white matter is 100%, means (circles) and medians (squares) are to the left with vibrissals showing ± 95% confidence intervals. Due to non-homogenous variance, Levene’s test p<0.001, Welch test used for comparison of the means did not show any significant difference, F(2,104.52)=1.05,p=0.36 with Games-Howell post-hoc n.s., between the reciprocally connected pairs (pink), unidirectionally connected pairs (powder blue) and non-connected pairs (baby blue). **E.** Euclidian distance between all the cell pairs were recorded within 120 µm of each other and show no difference, although the tendency of connected pairs having a smaller Euclidian intersomatic distance is showing (averageEuclidian distance between reciprocally connected pairs = 23.65 µm, median = 21.29 µm, n=18; between unidirectionally connected pairs avg = 26.96 µm, median = 24.22 µm, n=58; between non-connected pairs avg = 31.11 µm, median = 27.6 µm, n=119, One-Way ANOVA F=2.52, p=0.08 n.s.). Scale bar for A. is 50 µm.

This percentage falls within the broad range of previous reports on connectivity between those types of cells (Supp. Table 2) (Hage et al., 2022; Campagnola et al., 2022; Frandolig et al., 2019; Gainey et al., 2018; Espinoza et al., 2018; Jouhanneau et al., 2018; Guan et al., 2017; Jiang et al., 2015; Avermann et al., 2012; Hofer et al., 2011; House et al., 2011; Packer and Yuste, 2011; Gibson et al., 2008; Kapfer et al., 2007; Gabernet et al., 2005; Yoshimura and Callaway, 2005; Beierlein et al., 2003; Beierlein and Connors, 2002; Holmgren et al., 2003; Thomson and Morris, 2002; reviewed in Thomson and Lamy, 2007; Gupta et al., 2000; Ali et al., 1999). PV+ connectivity to Pyr neurons was present in 71 pairs (53 unidirectional and 18 bidirectional). Pyr connectivity to Pyr neurons was less frequent, occurring in only 24 pairs (6 unidirectional and 18 bidirectional). The relatively high connection probability from PV+ to Pyr neurons with reciprocal connectivity is striking (75%, 18 of 24, Fisher’s Exact 20.81, p=1.6E-5) (Yoshimura and Callaway, 2005). All the pairs were recorded within 120 µm of Euclidian distance apart (Fig. 4E). No significant difference was found within this distance between reciprocally connected, unidirectionally connected and non-connected pairs (average Euclidian distance between bidirectionally connected pairs = 23.65 µm, median = 21.29 µm; unidirectionally connected pairs avg = 26.96 µm, median = 24.22 µm; non-connected pairs avg = 31.11 µm, median = 27.6 µm; One-Way ANOVA F(2,192)=2.52, p= 0.08, Tukey post-hoc ns.; Fig 4E). Additionally, since connection probability between recorded pairs decreased with increasing distance (Hage et al., 2022; Packer and Yuste, 2011), we compared Euclidian distance with one-tailed independent samples Student’s t-test, due to smaller sample size for bidirectionally connected pairs (n=18) and known predicted difference of larger distance between non-connected pairs (n=119). The difference between bidirectionally connected pairs to non-connected pairs produced a statistically significant result (t-test(135) = 1.74, p=0.04). The recorded pairs did not differ in their distance to pia (due to non-homogenous variance, Levene’s test p<0.001, Welch test was used for comparison of the means did not show any significant difference, F(2,104.52)=1.045, p=0.36 with Games-Howell post-hoc ns.).

### Somatosensory cortical and thalamic excitation in connected vs. non-connected PV+ and Pyr neurons

To test whether excitation from somatosensory cortex and thalamus is organized differently in connected versus non-connected pairs of PV+ and Pyr cells, we compared whole-cell voltage-clamp responses to vS1 and PO excitation in simultaneously recorded pairs of neurons. The similar distribution of ChR2-induced EPSC amplitude with ReaChR induced EPSC amplitude (Fig. 3D) following the same amount of viral volume injected into PO and vS1 suggest both pathways excite M1 with roughly similar strength. To compare the responses across different slices and animals and account for the difference in ChR2 and ReaChR expression levels between individual animals, opsin-mediated EPSCs amplitudes were normalized to the maximum response in the slice during the 250 ms delay sweep (Fig. 5, Supp. Fig. 2; Supp. Fig. 3). Only the slices with at least 2 cells were included. To visualize both somatosensory cortical and thalamic inputs in the same pairs in the same graphs, we used scatter bubble plots, that allows to show the third dimension by controlling the size of the data points. Thus, plots comparing vS1 input to PV+ and Pyr neurons could also show the strength of PO input with the size of the marker. Visual inspection shows a difference in normalized EPSCs amplitudes between connected and non-connected pairs of PV+ and Pyr cells, which is emphasized when the thalamic inputs to PV+ or Pyr cells are chosen as a third dimension/variable (controlling the size of each data point in the pair receiving input from somatosensory cortex) (Fig. 5). Specifically, PV+ neurons receive stronger vS1 inputs than nearby Pyr neurons (Fig. 5A-B), resulting in most points falling below the unity line (gray). Furthermore, the scatter of these points is reduced in connected versus non-connected pairs (Fig 5A-F), resulting in fewer points scattered above the line, suggesting less variance. This trend also seems to hold for PO inputs to PV+ and Pyr neurons (Fig. 5G-L), though less pronounced. The normalized input strength is comparable for PV+ and Pyr in both groups, yet the distribution is shifted (Fig. 5O-P).

**Figure 5.**
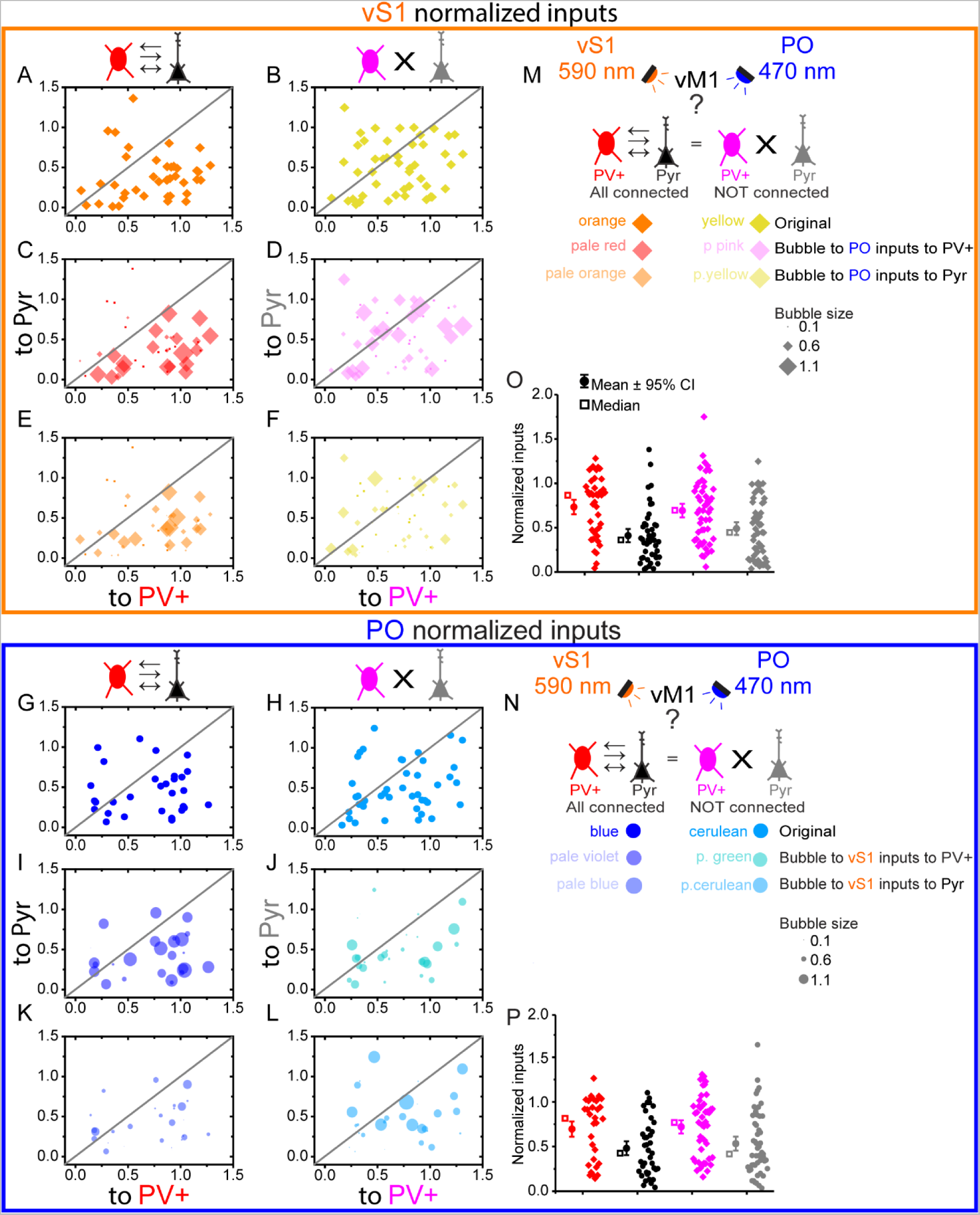
Normalized vS1 and PO inputs to connected and not pairs of PV+ and Pyr neurons show different trends. All inputs are from the 100 ms delay protocol normalized to the mazimum slice peak at 250 ms delay protocol. **A.C.E.** vS1 (orange) inputs for connected PV+ (red) and Pyr (black) pairs. **A.** Original figure. **C.** is A. bubble plotted to PO (blue) inputs to PV+ neurons in the same pairs. **E.** is A bubble plotted to PO inputs to Pyr in the same pairs. B.D.F. vS1 inputs for non-connected PV+ (magenta) and Pyr (gray) pairs. **B.** Original figure. **D.** is B. bubble plotted to PO inputs to PV+ neurons in the same pairs. **F.** is B. bubble plotted to PO inputs to Pyr in the same pairs. **G.I.K.** PO inputs for connected PV+ and Pyr pairs. **G.** Original figure. **I.** is G. bubble plotted to vS1 inputs to PV+ neurons in the same pairs. **K.** is G. bubble plotted to vS1 inputs to Pyr neurons in the same pairs. **H.J.L.** PO inputs for non-connected PV+ and Pyr pairs. **H.** Original figure. **J.** is H. bubble plotted to vS1 inputs to PV+ neurons in the same pairs. **L.** is H. bubble plotted to vS1 inputs to Pyr neurons in the same pairs. Bubble plots scale is from 0.1 to 1.6 with Δ of 0.5 **M.** Schematics of scientific inquiry question, methods to study it, bubble pltos color coding and size for vS1 inputs. **N.** Schematics of scientific inquiry question, methods to study it, bubble pltos color coding and size for PO inputs. **O.** is A. and B. shown as a box plots. **P.** is G. and H. shown as a box plots.

We then sought to test whether the connected cell pairs had correlated inputs. We speculated that interconnected subnetworks of neurons performing similar computations might get correlated input, such as both PV+ and Pyr neurons receiving strong or weak input. The alternative is that long-range input strength would be random with respect to whether pairs were connected. We plotted the EPSC amplitudes for connected and non-connected pairs (Fig. 6B,C,F,G). We then fit these with a linear regression, finding that EPSCs from both vS1 and PO inputs before normalization showed higher correlation in connected (vS1 inputs in connected pairs, adjusted to the degrees of freedom R^2^ = 0.21; PO inputs in connected pairs, R^2^=0.37) compared to non-connected pairs (vS1 inputs in non-connected pairs, R^2^ = 0.18; PO inputs in non-connected pairs, R^2^=0.13; Fig. 6, Supp. Fig. 4, and Supp. Fig. 5, the data was resampled 1000 times bootstrapped and produced correlation coefficients that were compared between those groups, p<E-10) suggesting that co-targeting of long-range projections is dependent on whether they contact connected or non-connected pairs.

**Figure 6.**
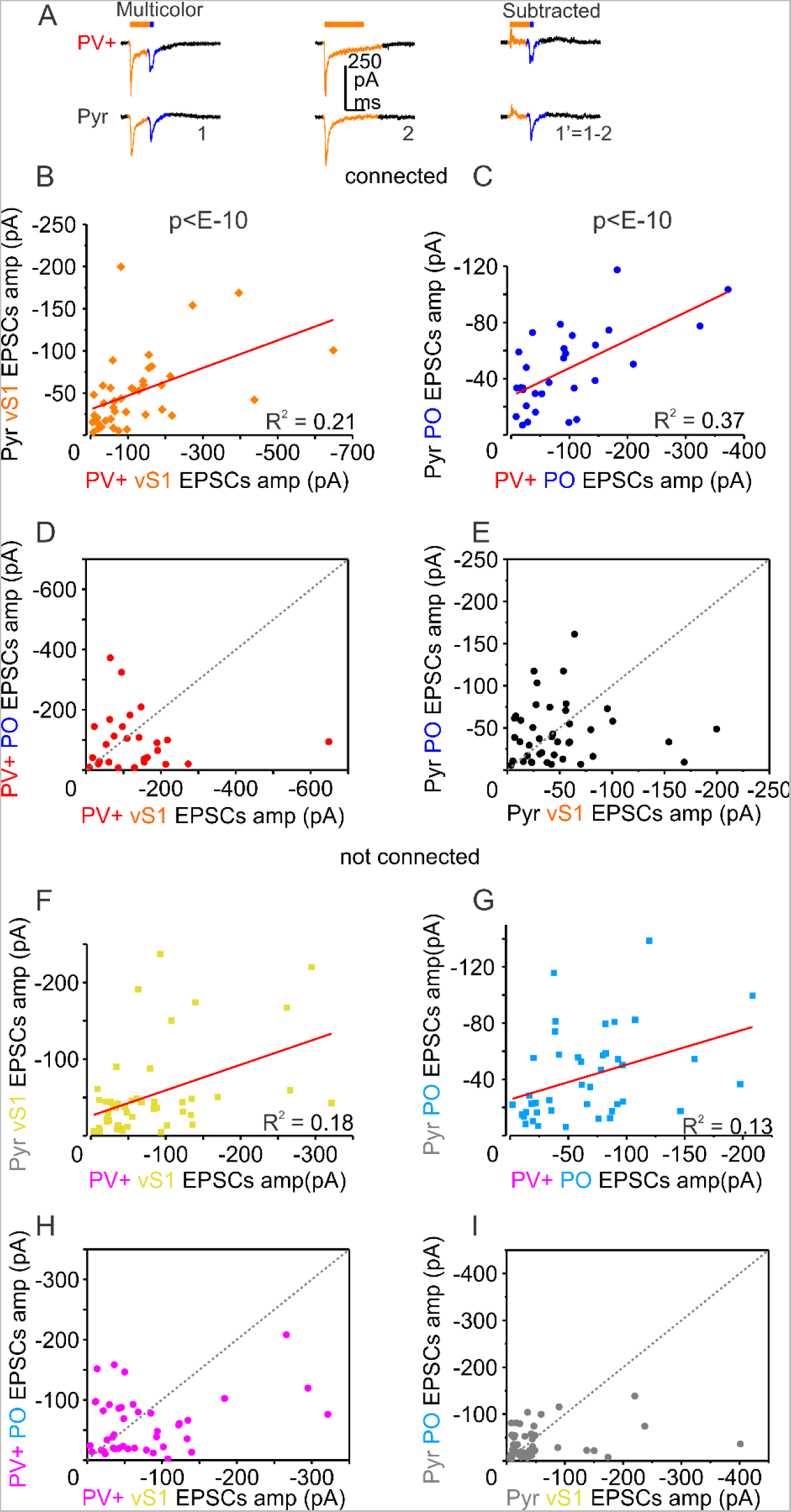
Increased correlation of the long-range inputs to connected pairs. **A.** Example of voltage-clamp traces from PV+ (red) and Pyr pair with 590 nm stimulation of vS1 axons and 470 nm stimulation of PO (blue) axons middle traces are 590 nm stimulation alone and the right traces show the result of subtraction of middle traces from the first traces. Connected pairs are B.C.D.E. **B.** Scatter plot of vS1 in connected pairs showing a higher correlation compared to non-connected pairs in F. The data in B. and F. was resampled using 1000 samples bootstrap and produced Spearman ρ correlation coeficients, which were squared and compared with nonparametric independent samples Mann-Whitney test, U=208790.0, p<1E-10. **C.** Scatter plot of PO EPSCs in connected pairs showing a higher correlation compared to non-connected pairs in G. The data in C. and G. was resampled using 1000 samples bootstrap and produced Spearman ρ correlation coeficients, which were squared and compared with nonparametric independent samples Mann-Whitney test, U=279207.0, p<1E-10. **D.** Scatter plot of PO and vS1 EPSCs in the PV+ interneurons from the connected pairs. **E.** Scatter plot of PO and vS1 EPSCs in the Pyr neurons from connected pairs. Non-connected pairs are F.,G.,H.,I. **F.** Scatter plot of vS1 EPSCs from non-connected pairs (yellow). **G.** Scatter plot of PO (teal) EPSCs in non-connected pairs. **H.** Scatter plot of PO and vS1 EPSCs in the PV+ (magenta) interneurons from the non-connected pairs. I. Scatter plot of PO and vS1 EPSCs in the Pyr neurons from the non-connected pairs. Correlations are estimated by the adjusted to the degrees of freedom Pearson R^2^.

Normalized EPSCs showed that vS1 preferentially targets PV+ compared to Pyr neurons regardless of whether Pyr-PV+ pairs were connected, but connected pairs showed a bigger difference between the Pyr EPSCs (connected Pyr normalized mean = 0.41; non-connected Pyr normalized mean= 0.49) and PV+ EPSCs (connected PV+ normalized mean =0.73; non-connected PV+ normalized mean= 0.69), mean difference (connected Δ =0.32; non-connected Δ= 0.2) with larger variance in non-connected pairs (connected PV+ STDEV^2^= 0.11, connected Pyr STDEV^2^=0.09; non-connected PV+ STDEV^2^= 0.13, non-connected Pyr STDEV^2^= 0.11) (Fig. 5O). The PO inputs were larger in the same pairs with PV+ preference in connected cells, while large PO inputs were more broadly distributed in non-connected cells (Fig. 5C, D).

### Excitation to Inhibition ratio of vS1 and PO inputs

We wanted to test how the recruitment of PV+ cells by long-range projections is correlated with the ReaChR and ChR2 induced IPSCs in both PV+ and Pyr cells. As before, we thought that interconnected subnetworks of neurons might get correlated excitatory and inhibitory input. Thus, we voltage clamped the pairs at 0 mV after acquiring the excitatory responses. For both PO and vS1 inputs, opsin-induced IPSCs correlation with EPSCs in L2/3 and L5A PV+ and Pyr cells was assessed with adjusted to the degrees of freedom Pearson R^2^. Correlation of vS1-evoked IPSCs to EPSCs in L2/3 PV+ neurons was R^2^=0.44 (Fig. 7B). In L5A PV+ neurons, it was R^2^=0.59. For L2/3 Pyr cells, vS1-evoked IPSCs to EPSCs correlation was R^2^=0.49, and, in L5A Pyr cells, it was R^2^=0.72. The PO-evoked IPSCs to EPSCs correlation in L2/3 PV+ cells was R^2^=0.71, and, in L5A PV+ cells, it was R^2^=0.34. PO-evoked IPSCs to EPSCs correlation in L2/3 Pyr cells was R^2^=0.42, with Pyr cells having less correlation from PO inputs in L5A (R^2^=0.04). L2/3 Pyr cells also had an increased IPSCs amplitudes in L2/3 from vS1 inputs (Fig. 7B, 7C). There was also less correlation of IPSCs to EPSCs in non-connected Pyr cells from both vS1 and PO inputs (non-connected Pyr vS1 EPSCs to IPSCs R^2^=0.03; non-connected Pyr PO EPSCs to IPSCs R^2^=0.07, Fig. 8A,8B).

**Figure 7.**
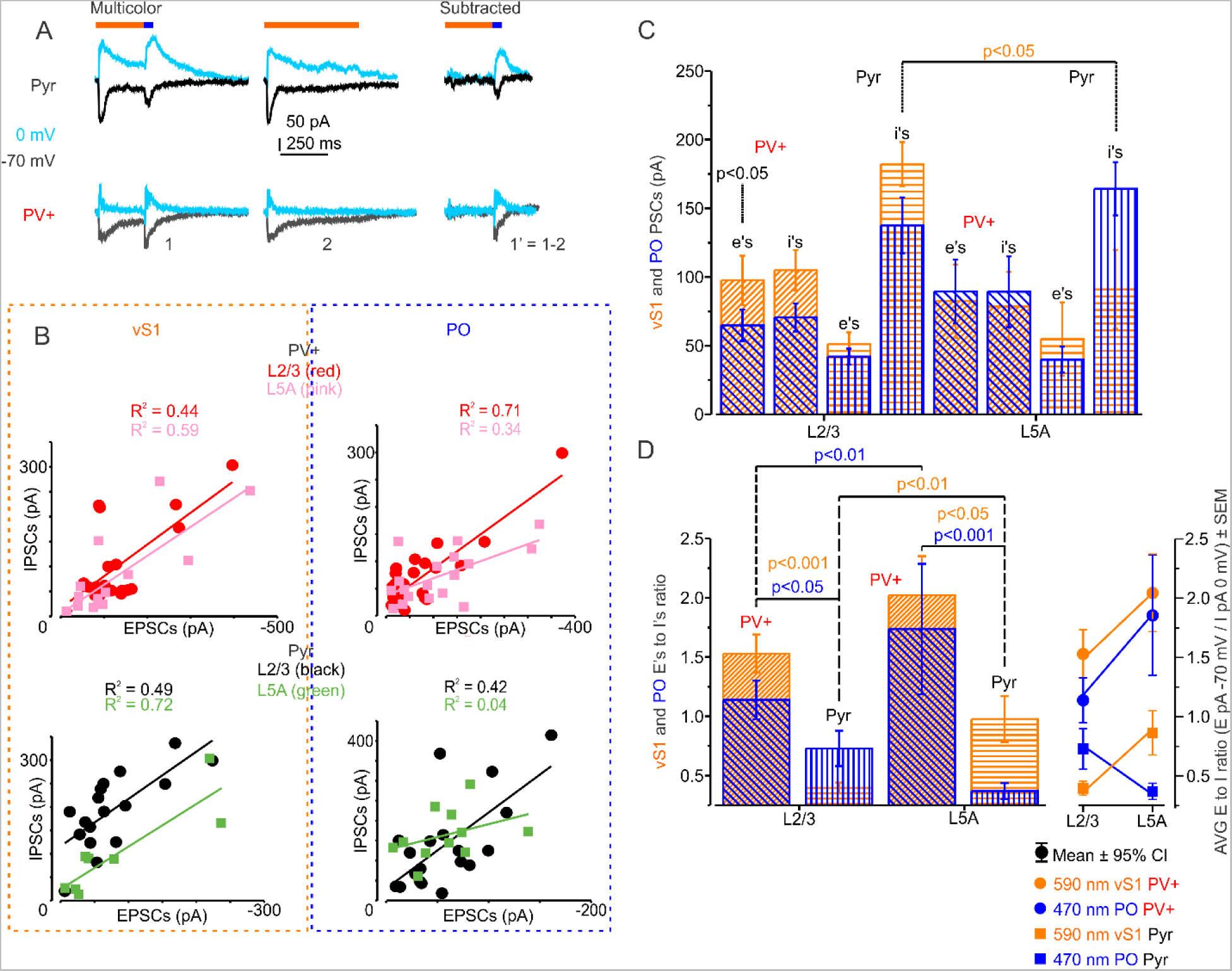
Long-range inputs excite more PV+ interneurons eliciting stronger feedforward inhibition in layer-specific manner. **A**. Example traces showing a pair of PV+ and Pyr neurons recorded at -70 mV and at 0 mV (teal) holding potential. **B.** Scatter plot of EPSCs and IPSCs for vS1 inputs (upper panel), and PO (lower panel) for PV+ (left panel) and Pyr (right panel). Cortical layers 2/3 data for PV+ (red) and L5A (pink), for Pyr the L2/3 data is in black and L5A is in brown. **C.** vS1 and PO EPSCs (e’s) at -70 mV holding potential multiplied by -1 for the convinience of presentation, and IPSCs (I’s) at 0 mV holding potential; vS1 I’s wre significantly larger in L2/3 than L5A Pyr (independent samples Mann-Whitney (M-W) U=39, p=3.08E-2). vS1 inputs were significantly larger than PO inputs to L2/3 PV+ (M-W U=2308, p=1.97E-2). **D.** Averaged layer-specific Excitation to Inhibition ratio where the amplitude of EPSCs at -70 mV is divided by the amplitude of IPSCs at 0 mV for each cell. E to I ratio for vS1 and PO inputs was significantly larger in PV+ than in Pyr in L2/3 (M-W U=52 and 200.5 (PO), p=2.2E-5 and 4.66E-2 (PO)); vS1 inputs E to I ratio was significantly larger in L5A than L2/3 Pyr (M-W U=182, p=4.89E-3); PO inputs E to I ratio was significantly larger in L5A than in L2/3 PV+ (M-W U=62.5, p=2.09E-3); vS1 and PO inputs E to I ratio was significantly larger in L5A PV+ than Pyr (M-W U=49 and 41.5 (PO), p=1.83E-2 and 7.6E-5 (PO)). Correlation is estimated by the adjusted by the degrees of freedom Pearson R^2^.

**Figure 8.**
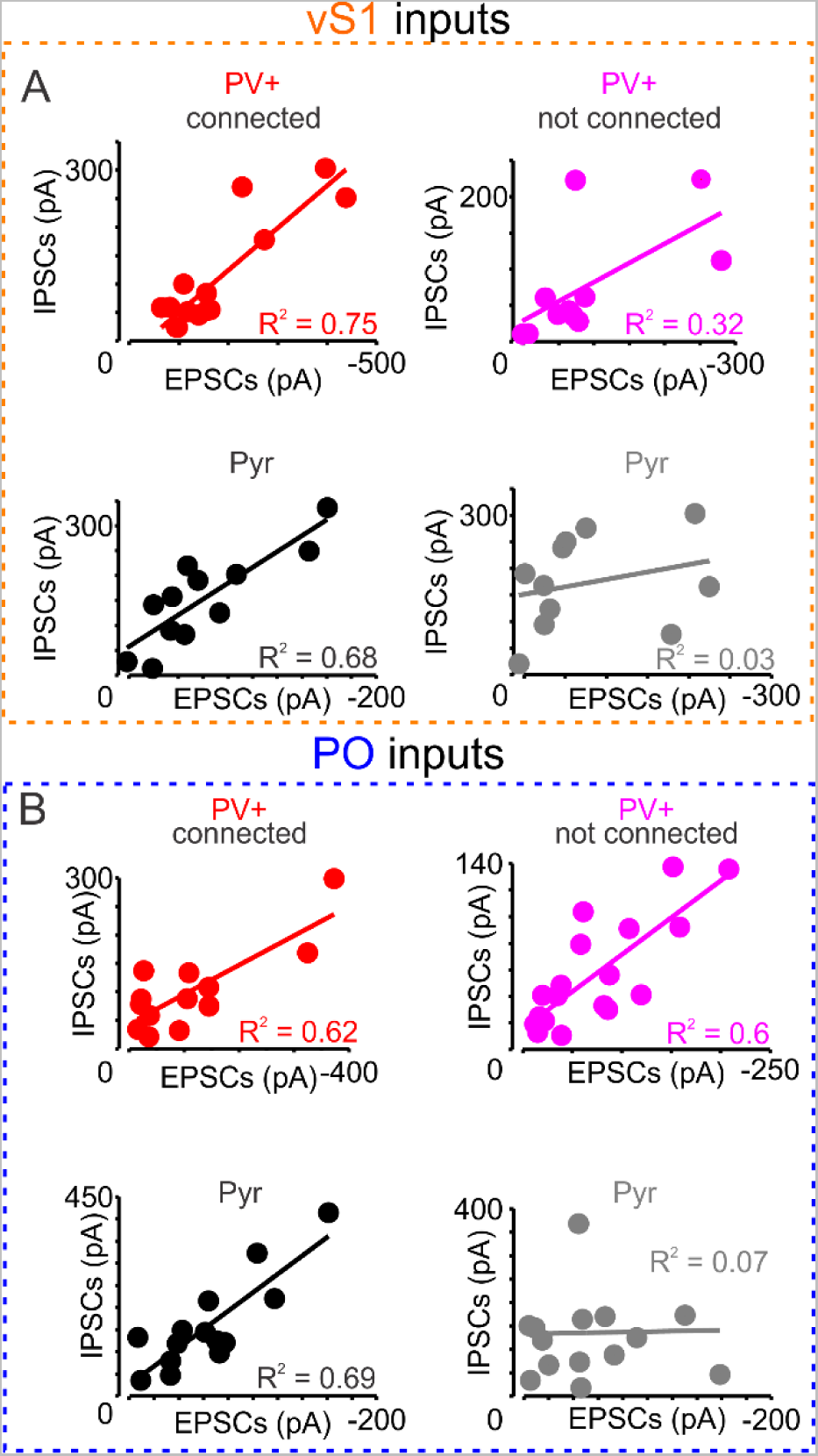
Correlation of long-range excitatory and feedforward inhibitory inputs is weaker in non-connected Pyr cells. **A.** Scatter plots of vS1 (orange), and **B.** PO (blue) inputs EPSCs (x axis) and IPSCs (y axis) for connected (left panels) PV+ (red) and Pyr (black) and non-connected (right panels, magenta for PV+ and gray for Pyr) .

Comparison of excitatory to inhibitory response ratio within the same cells between L2/3 and L5A showed that PO excitatory drive is stronger in PV+ cells in L5A compared to L2/3 PV+ cells (Mann-Whitney, p=2.09E-3; Fig. 7C,D), confirming the results from our previous studies with sCRACM (Okoro et al. 2022). For Pyr cells the vS1 excitatory drive was larger in L5A compared to L2/3 (E to I ratio, Mann-Whitney, p=4.89E-3; Fig. 7C,D) confirming the results from a previous study (Mao et al. 2011). For both L2/3 and L5A, the vS1 and PO excitatory drive was larger for PV+ cells than for Pyr cells (L2/3 vS1 inputs E/I ratio Mann-Whitney, p=2.2E-5; PO inputs E/I ratio Mann-Whitney, p=4.66E-2; L5A vS1 inputs E/I ratio Mann-Whitney, p=1.83E-2; L5A PO inputs E/I ratio Mann-Whitney, p=7.6E-4; Fig. 7C,D). Consistent with larger vS1 inputs in L2/3 PV+ cells the vS1 inhibitory responses were larger in L2/3 Pyr cells compared to L5A Pyr cells (Mann-Whitney, p=3.08E-2; Fig. 7B,7C,7D). This suggests that long-range inputs excite more PV+ cells eliciting stronger feedforward inhibition in a layer-specific manner.

## Discussion

Excitatory subnetworks are a set of connected Pyr cells receiving shared interlaminar (excitatory), intralaminar (inhibitory and excitatory) or long-range inputs (Yoshimura et al., 2005; Morgenstern et al., 2016b), or sharing a single inhibitory cell as a hub (Palagina et al., 2019; Yoshimura and Callaway, 2005). Local connectivity supports the existence of such excitatory subnetworks within small anatomical loci (Faber et al., 2019; Palagina et al., 2019; Vegué et al., 2017; Lee, W. A. et al., 2016; Yoshimura et al., 2005). Further, subnetworks might share common functional properties, such as receptive fields in visual areas (Ko et al., 2011; Ohki et al., 2006; 2005).

However, the degree to which interneurons participate in such subnetworks is unclear. Imaging interneuron response properties in visual cortex suggests that these cells are more broadly tuned than excitatory neurons (Kerlin et al., 2010), while other studies suggest PV+ cells may be selective to orientation and direction (Runyan et al., 2010). Further, in direction- or orientation-selective inhibitory interneurons (52%) and their clusters, 75% of clustered Pyr cells shared direction tuning with their corresponding inhibitory neuron (Palagina et al., 2019). Relatively dense local connectivity of PV+ neurons has been proposed, with both nonspecific but frequent output to local Pyr neurons (Packer and Yuste, 2011; Fino et al., 2013) and pooled excitatory input from Pyr cells with different properties (Sohya et al., 2007). How to reconcile a role for selective inhibitory connectivity with non-selective all-to-all inhibition remains to be answered. One possibility is that connections are common, but synaptic strength selectively varies within or across subnetworks (Znamenskiy et al., 2018). One statistical/structural explanation for subnetworks is the targeting by feedforward and feedback long-range projections, which may increase the information propagation and processing in cortical circuits by selecting clustered cells that have higher probability of connecting to each other (Peron et al., 2020; Faber et al., 2019; Palagina et al., 2019; Rost et al., 2018; Nigam et al., 2016). Thus, synaptic inputs from long-range projections may be correlated in connected clusters.

Motor cortex is somatotopically organized, and thus might contain distinct subnetworks for different movements of the same limb. Thus, we tested the local and long-range connectivity of PV+ and Pyr neurons to assess the existence of specific subnetworks. Here, we show that connected pairs of PV+ and Pyr cells in mouse motor cortex (vM1) have stronger correlation of long-range somatosensory (vS1) and thalamic (PO) projections than non-connected pairs (Fig. 6). Thus, subnetworks with strong vS1 input exist and more strongly excite connected PV+ and Pyr neurons. Further, the contribution of interneurons to subnetworks is not simply correlated connection strength, but also enhanced connection probability. While PV+ neurons make frequent local connections to Pyr neurons (N=71/197 pairs, 36.0%), Pyr neurons make sparser connections to PV+ cells (N=24/197, 12.2%). However, Pyr neurons are much more likely to excite PV+ cells that reciprocally connect to them, with connection rates as high as those for PV+ output (N=18/53, 34.0%). Thus, connected pairs were reciprocally connected at much higher than random rate (Fig. 4). The overall connection probability is potentially underestimated, as some connections may be severed in the brain slices. But it is unlikely that slice preparation differentially severs connections between neurons that lack correlated excitatory input.

This data shows that, consistent with earlier work (Hooks et al., 2015), single Pyr and PV+ neurons receive input from both cortical (vS1) and thalamic (PO) sources. In assessing strength of connections, vS1 inputs were stronger in L2/3 PV+ cells than PO inputs (Fig. 3). Furthermore, both classes of excitatory inputs were much stronger in PV+ cells than to Pyr cells (Fig. 3, Fig. 5). This is in line with previous studies showing increased thalamocortical inputs to PV+ compared to Pyr cells in somatosensory cortex (Cruikshank et al., 2007; 2010; Gabernet et al., 2005; Gibson, J. R. et al., 1999). There may be differences in measured EPSC strength due to differences between the cell types. PV+ cells may be more electrotonically compact, making larger inputs easier to measure, and differences in intrinsic excitability may favor higher firing rates in PV+ neurons (Supp. Fig. 1). These differences amplify the effectiveness of inputs in PV+ neurons. Thus, in general, the E/I ratio of PV+ cells is higher than the E/I ratio of Pyr cells (Fig. 7). Further, comparing input strength across layers (Fig. 7D), E/I ratio for vS1 input to Pyr cells increases from L2/3 to L5A implicating greater recruitment of feedforward inhibition in L2/3, presumed to originate from PV+ cells. In contrast, E/I ratio for thalamic input to Pyr cells goes down from L2/3 to L5A, implicating greater recruitment of feedforward inhibition in L5A (Okoro et al., 2022).

This work proposes a role for long-range projections as part of the neural circuit organization that differentiates the cortex into subnetworks. Differences in cortical and thalamic input to different local subnetworks can result in local circuit elements specialized for processing different streams of information. However, these experiments are done *ex-vivo* in mouse cortical slices, which may underestimate connectivity. Whether similar results can be obtained *in-vivo* is still remains to be tested.

## Acknowledgements

We thank Y Kate Hong, Qian-Quan Sun, Alison Barth, and other members of the Hooks lab for comments and suggestions. We also thank Quincy Erickson-Oberg for helping with tracing biocytin filled neurons. This work was supported by a CDMRP PRMRP Discovery Award PR201842 (RUG and BMH), a NARSAD Young Investigator Award (BMH), and NIH NINDS R01 NS103993 (BMH).

## FIGURE LEGENDS

### SUPPLEMENTARY MATERIAL

**Supplementary figure 1.**
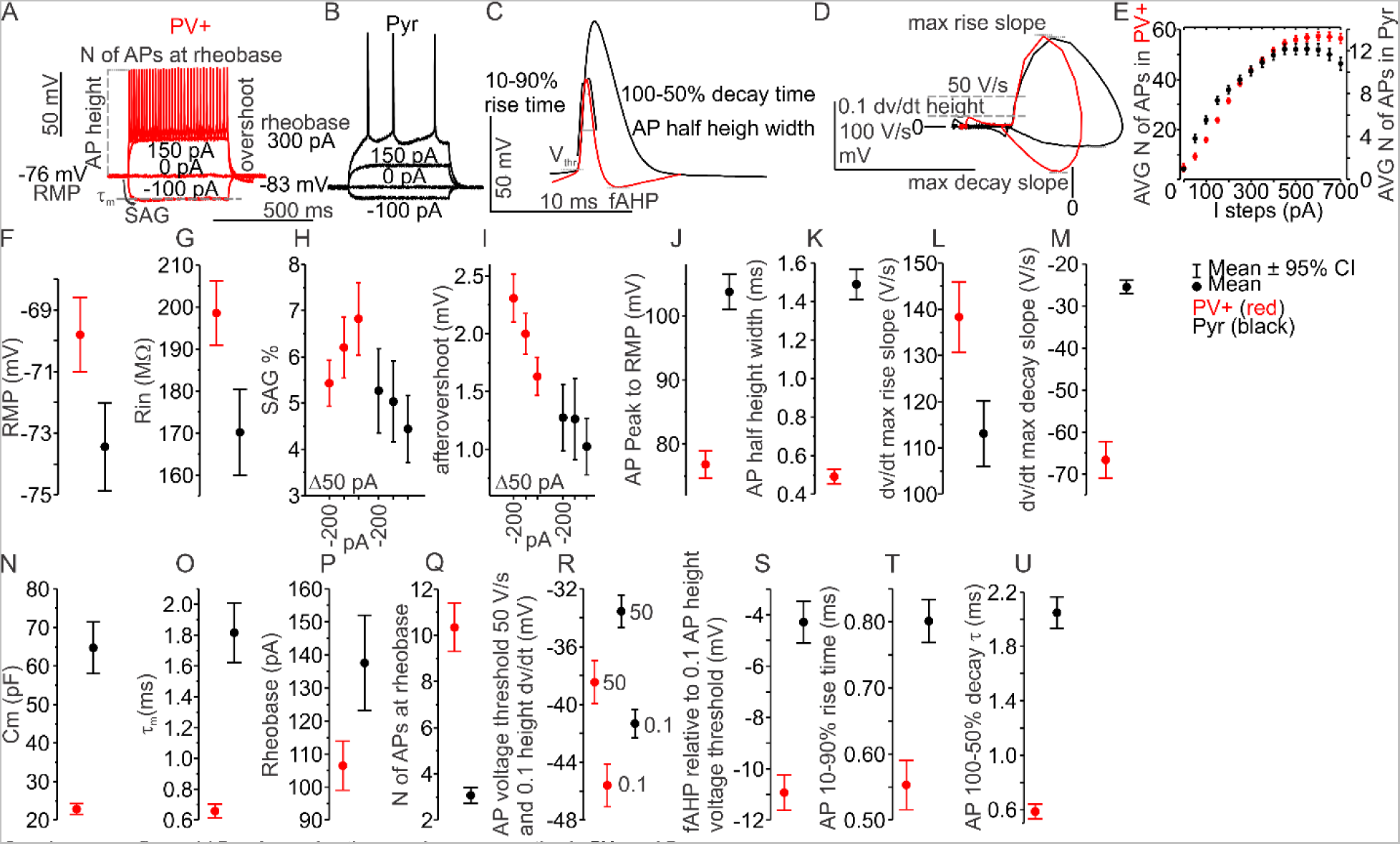
Passive and active membrane properties in PV+ and Pyr neurons. **A.** Example current-clamp traces from PV+ (red) with current steps shown in between, resting membrane potential (RMP) to the left measured before current step application, SAG is the negative deflection below resting state to the negative current steps in percent; Action Potential (AP) height is measured from RMP; the rheobase is the minimal voltage required to make the neuron fire AP (300 pA); number of APs at rheobase is also recorded; overshoot is measured at the end of 500 ms negative current steps. **B.** Example current-clamp traces from Pyr (rheobase 300 pA). **C.** Example APs from PV+ and Pyr showing 10-90% rise time, voltage threshold (Vthr), fast AfterHyperPolarization (fAHP); 100-50% decay time; and AP half-height width. **D.** The APs from C. are converted to phase-space plot to show the 50 V/s voltage threshold, 0.1 dv/dt height voltage threshold; max rise slope; max decay slope. **E.** Input-output curve showing number of APs (average ± SEM) fired by the cells in response to the 500 ms current steps. **F.-U.** Membrane properties.

**Supplementary Figure 2.**
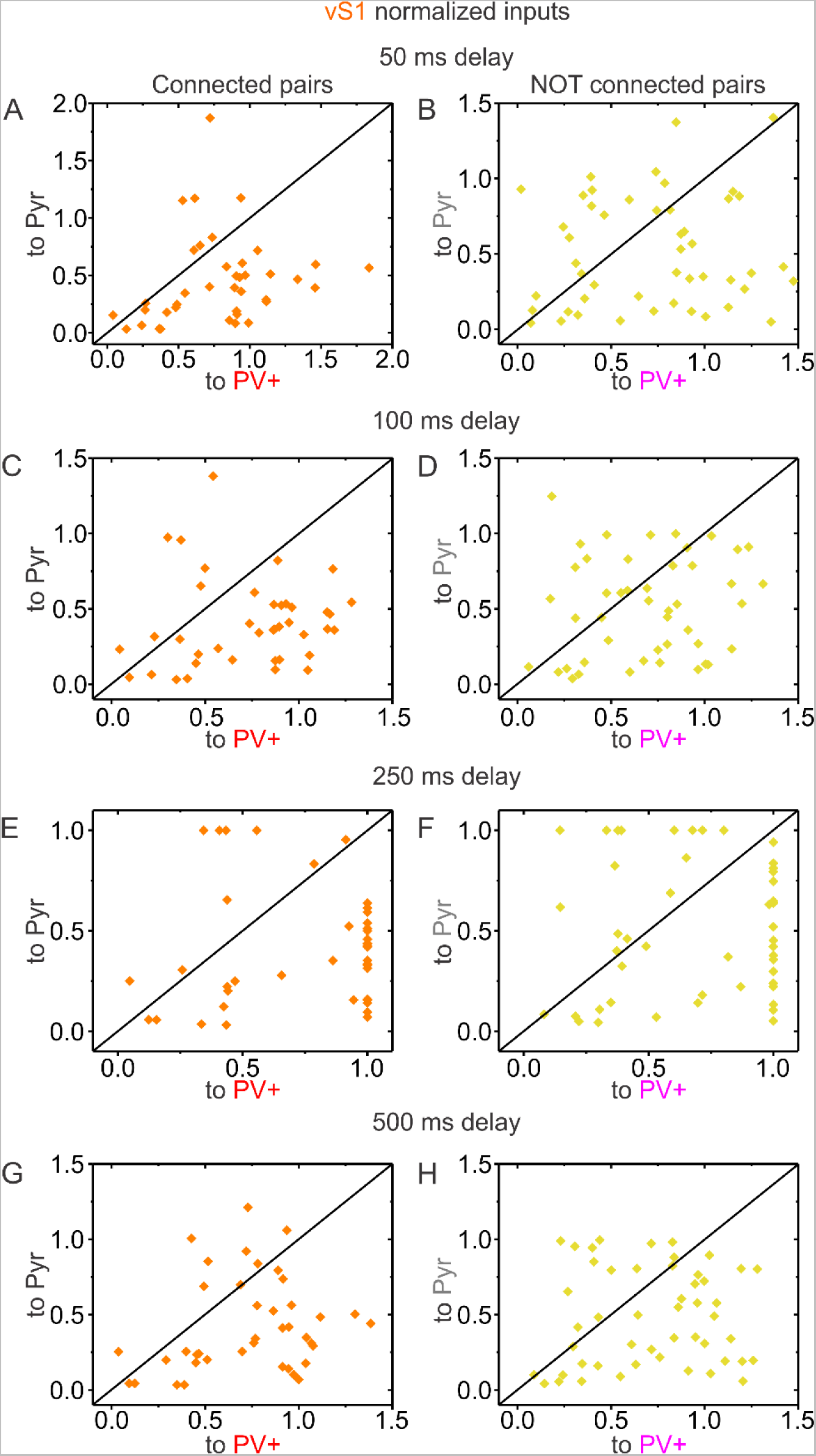
Normalized vS1 inputs EPSCs amplitude peaks to the slice maximum at 250 ms delay protocol. **A., C., E., G**. Connected PV+ (red) and Pyr (black) pairs. **B., D., H., F.** Non-connected PV+ (magenta) and Pyr (gray) pairs. **A., B.** 50 ms delays protocol **C., D.** 100 ms delay protocol. **E., F.** 250 ms delay protocol. **G., H.** 500 ms delay protocol.

**Supplementary Figure 3.**
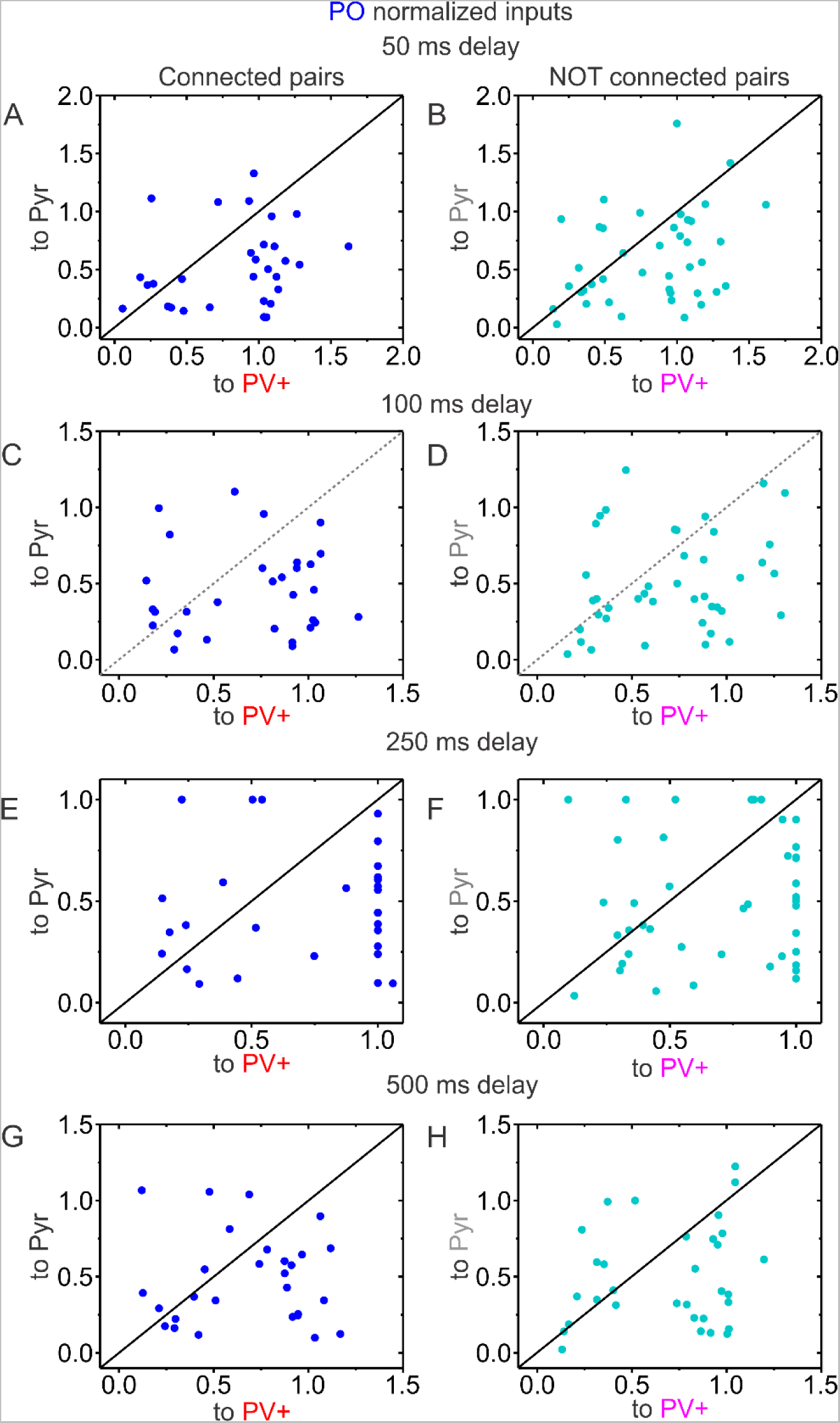
Normalized PO inputs EPSCs amplitude peaks to the slice maximum at 250 ms delay protocol. **A., C., E., G.** Connected PV+ (red) and Pyr (black) pairs. **B., D., H., F.** Non-connected PV+ (magenta) and Pyr (gray) pairs. **A., B.** 50 ms delays protocol **C., D.** 100 ms delay protocol. **E., F.** 250 ms delay protocol. **G., H.** 500 ms delay protocol.

**Supplementary Figure 4.**
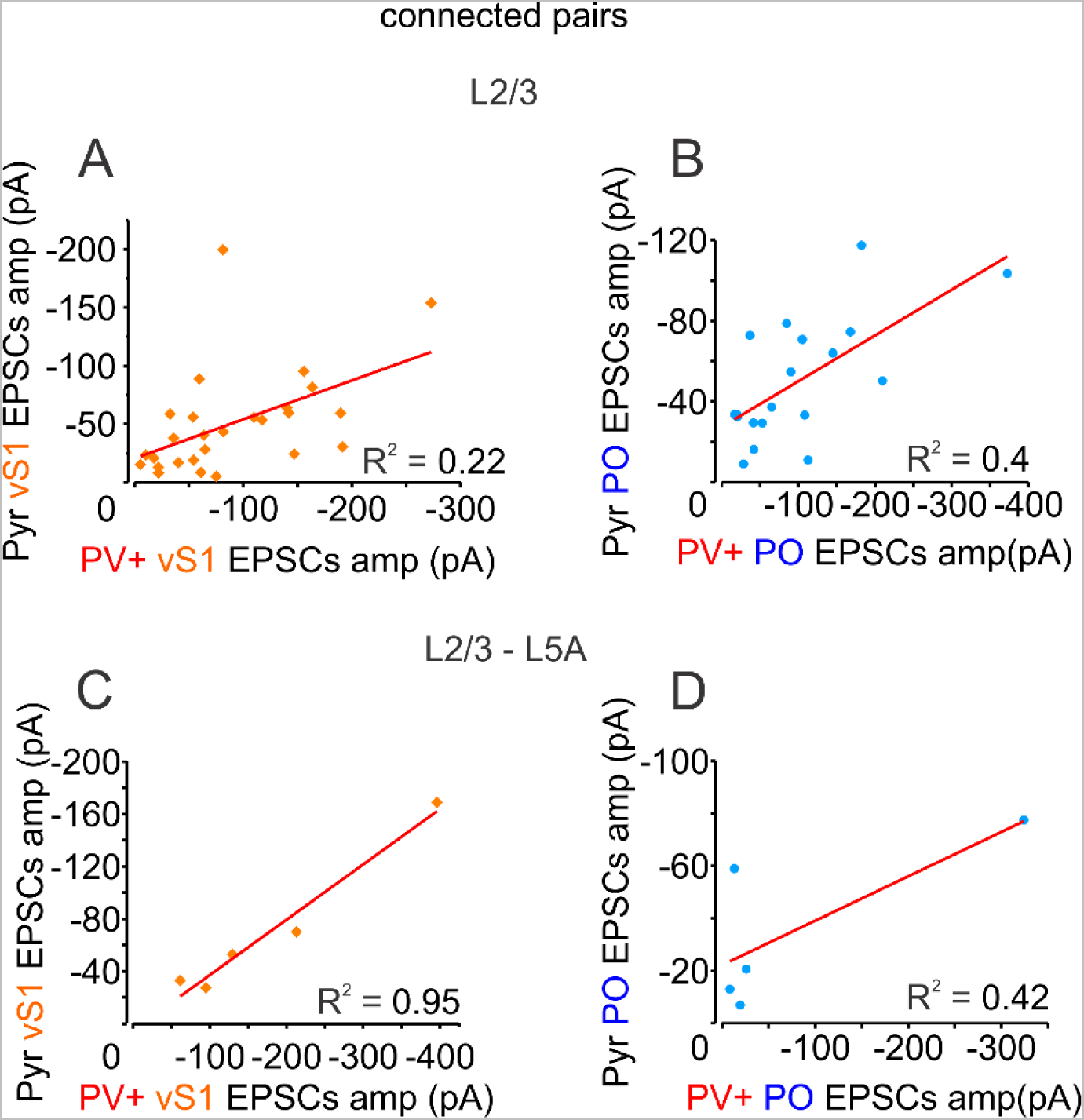
Long-range inputs have a layer-specific differential correlation for connected pairs. **A.** Scatterplot of vS1 EPSCs in connected pairs in layer 2/3. **B.** Scatter plot of PO EPSCs in connected pairs in layer 2/3. **C.** Scatter plot of vS1 EPSCs in connected pairs between layer 2/3 and 5A. **D.** Scatter plot of PO EPSCs in connected pairs between layer 2/3 and 5A.

**Supplementary Figure 5.**
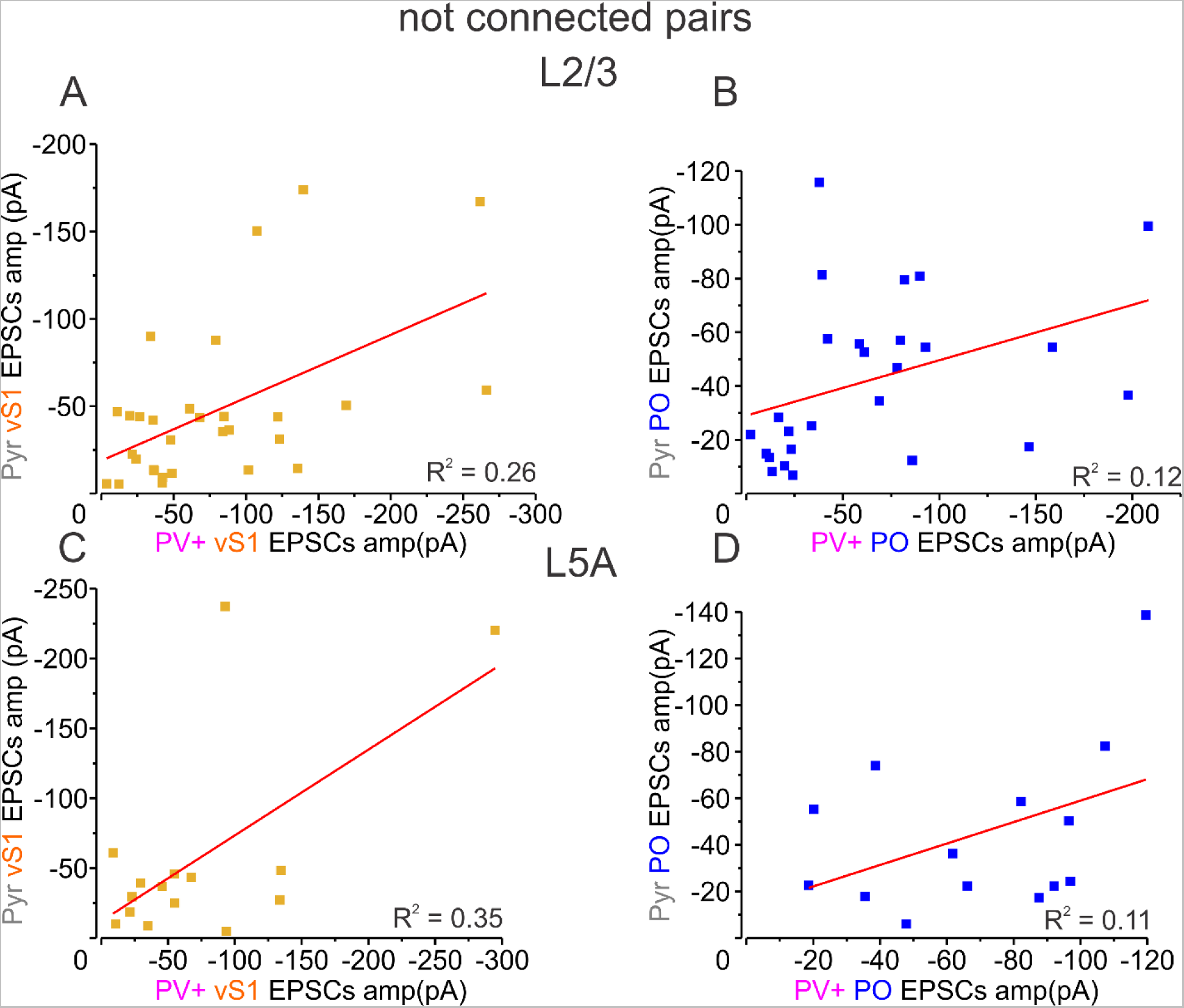
Long-range inputs have a layer-specific differential correlation for non-connected pairs. **A.** Scatter plot of vS1 EPSCs in non-connected pairs in layer 2/3. **B.** Scatterplot of PO EPSCs in non-connected pairs in layer 5A. **C.** Scatterplot of vS1 EPSCs in non-connected pairs in layer 5A. **D.** Scatterplot of PO EPSCs in non-connected pairs in layer 5A.

**Supplementary Table 1.** Intrinsic Cell Properties of PV+ and Pyr Neurons in M1.

|  |  | PV+ |  |  | Pyr |  |  |
| --- | --- | --- | --- | --- | --- | --- | --- |
|  |  | AVG | STDEV | N | AVG | STDEV | N |
| RMP | mV | -69.80 | 7.49 | 143 | -73.44 | 8.85 | 140 |
| Rin | MΩ | 209.97 | 90.56 | 147 | 172.66 | 140.10 | 145 |
| Cm | pF | 23.01 | 8.91 | 147 | 65.04 | 42.10 | 145 |
| taum | ms | 0.66 | 0.28 | 147 | 1.83 | 1.21 | 145 |
| Rin from V-I | MΩ | 197.54 | 47.30 | 140 | 169.65 | 64.26 | 136 |
| SAG -200 pA | % | 5.44 | 3.09 | 140 | 5.30 | 5.73 | 135 |
| SAG -150 pA | % | 6.21 | 4.12 | 139 | 5.06 | 5.49 | 136 |
| SAG -100 pA | % | 6.87 | 4.91 | 140 | 4.48 | 4.54 | 136 |
| After-Overshoot Peak at 1.5 s -200 pA | mV | 2.30 | 1.27 | 135 | 1.29 | 1.81 | 131 |
| After-Overshoot Peak at 1.5 s -150 pA | mV | 1.98 | 1.07 | 134 | 1.28 | 2.21 | 132 |
| After-Overshoot Peak at 1.5 s -100 pA | mV | 1.61 | 0.97 | 136 | 1.04 | 1.54 | 134 |
| Rheobase | pA | 106.52 | 46.34 | 138 | 138.52 | 90.38 | 135 |
| N of AP at rheobase | N | 10.29 | 6.48 | 140 | 3.10 | 2.13 | 136 |
| AP at rheobase voltage thr 50 V/s | mV | -38.50 | 9.29 | 137 | -33.51 | 7.10 | 131 |
| AP at rheobase voltage thr 0.1 height dV/dt V/s | mV | -45.64 | 9.22 | 138 | -41.29 | 6.17 | 136 |
| AP at rheobase Peak to RMP | mV | 76.89 | 13.42 | 138 | 103.63 | 17.00 | 136 |
| AP at rheobase trough relative to 0.1 height dV/dt V/s thr | mV | -10.98 | 4.37 | 79 | -4.32 | 5.20 | 104 |
| dV/dt Max rise slope | V/s | 138.55 | 47.46 | 138 | 113.03 | 44.56 | 136 |
| dV/dt Max decay slope | V/s | -66.82 | 27.29 | 138 | -25.39 | 9.90 | 136 |
| AP half height width | ms | 0.49 | 0.23 | 138 | 1.49 | 0.49 | 136 |
| AP 10-90% Rise Time | ms | 0.55 | 0.23 | 138 | 0.80 | 0.20 | 134 |
| AP 100-50% Decay Time | ms | 0.58 | 0.35 | 138 | 2.05 | 0.73 | 133 |

**Supplementary Table 2.** Comparison of PV+ and Pyr neuron pair recording data across the literature. Data for all pairs. The data are dominated by intralaminar connections, since PV+ to Pyr, or Pyr to PV+ interlaminar connections are more scarce. Paired or multi-paired: refers to whole-cell patch clamp of 2 or more neurons. Sharp: refers to sharp electrode intracellular recording. 2P: refers to two-photon stimulation and recording. ChR: refers to optogenetic expression of light sensitive actuator molecules (channelrhodopsin variants), through viral vector or genetic crossing. Rubi-glut: refers to Rubi-glutamate uncaging. Intersomatic distance is reported as a measure of Euclidian distance. Reported a horizontal offset if Euclidian distance was not calculated. In area column, S – somatosensory, V – visual, M – motor. Where no distinction between the interneuron types is made, both types are included in the percentage calculation. GC – granule cells. CThN – corticothalamic neurons. CCN – corticocortical neurons. Upper layer 6a – is the top 40% of the layer height, lower layer 6a - is the bottom 40% of the layer height. FS – fast-spiking interneurons, are considered to be a part of PV+ cells, LTS-low-threshold spiking interneurons, are considered to be a part of SOM+ cells. RS – regular spiking neurons, mostly correspond to Pyr cells. Cg1/2 – prefrontal cingulate cortex area 1/2 (dACC – dorsal anterior cingulate cortex). Depressing synapses are presumed to be from PV+ interneurons.

| Report + recording method | Total (n) | Connected (n) | % | Species | Area, Layers | Age | Intersomatic distance |
| --- | --- | --- | --- | --- | --- | --- | --- |
| Current work (paired) | | | | mouse | vM1, L2/3-5b | P28-123, AVG 46 | <120 μm |
| Pyr → PV+ | 197 | 6 | 3.05 | | | | |
| PV+ → Pyr | 197 | 53 | 26.9 | | | | |
| PV+ ↔ Pyr | 197 | 18 | 9.14 | | | | |
| Hage et al. 2022<br>(paired + 2P ChR) |  |  |  | mouse | V1, L2/3-6 | P40-126, AVG<br>58 | <100 μm |
| Pyr → PV+ | 28 | 13 | 46 |  |  |  |  |
| PV+ → Pyr | 156,31 | 35,17 | 22.4(2P)-<br>55 |  |  |  | <200(2P) μm |
| PV+ ↔Pyr |  |  | missing |  |  |  |  |
| Jiang et al. 2015<br>(multi-paired) |  |  |  | mouse | V1, L2/3-5 | ≥~P60 | <50 μm |
| Pyr → PV+ | 20(L5)-<br>82(L2/3) | 3(L5)-<br>13(L2/3) | 15(L5)-<br>15.9(L2/3<br>) |  |  |  |  |
| PV+ → Pyr | 20(L5)-<br>82(L2/3) | 5(L5)-<br>27(L2/3) | 25(L5)-<br>32.9(L2/3<br>) |  |  |  |  |
| PV+ ↔Pyr | 20(L5)-<br>82(L2/3) | estimate<br>based on<br>connected<br>Pyr (3-13) | ≤15 |  |  |  |  |
| Yoshimura & Callaway<br>2005 (paired) |  |  |  | rat | V1, L2/3 | P21-26 | <100 μm |
| Pyr → PV+ | 43 | 1 | 2.33 |  |  |  |  |
| PV+ → Pyr | 43 | 13 | 30.23 |  |  |  |  |
| PV+ ↔Pyr | 43 | 7 | 16.28 |  |  |  |  |
| Thomson et al. 2002<br>(multi-paired sharp) |  |  |  | cat | V1, L2/3-5 | not reported<br>(young adult) | missing |
| Pyr → PV+ & SOM+ | 25 | 2 | 8(L2/3) |  |  |  |  |
| PV+ & SOM+ → Pyr | 25 | 4 | 16(L2/3) |  |  |  |  |
| PV+ & SOM+ ↔Pyr | 25 | 3 | 12(L2/3) |  |  |  |  |
|  |  |  |  | rat | S, M, V,<br>L2/3-5 | not reported<br>(young adult) |  |
| Pyr → PV+ & SOM+ | 107 | 19 | 17.76(L2/<br>3) |  |  |  |  |
| PV+ & SOM+ → Pyr | 107 | 14 | 13.08(L2/<br>3) |  |  |  |  |
| PV+ & SOM+ ↔Pyr | 107 | 3 | 2.8(L2/3) |  |  |  |  |
| Beierlein et al. 2003<br>(paired) |  |  |  | rat | S, L4 | P14-21 | <50 µm |
| Pyr → PV+ | 172 | 74 | 43 |  |  |  |  |
| PV+ → Pyr | 190 | 83 | 44 |  |  |  |  |
| PV+ ↔Pyr | 190 | 34 | 17.89 |  |  |  |  |
| Packer and Yuste 2011<br>(paired &2P + Rubi-glut) |  |  |  | mouse | S, L2/3, L5,<br>frontal L2/3 | P12-45 | <200 µm |
| Pyr → PV+ | not<br>tested | not tested | not<br>tested |  |  |  |  |
| PV+ → Pyr | 27 | 24 | 88.89 |  | S,L2/3 | P13-16 | <100 μm |
|  | 7 | 6 | 85.71 |  | S,L5 |  |  |
|  | 5 | 4 | 80 |  | F, L2/3 |  |  |
|  | Fig. 4,5 | Fig. 4,5 | 71(2P) |  | S,L2/3 |  |  |
|  | Fig. 4,5 | Fig. 4,5 | 92(2P) |  | S,L5 |  |  |
|  | Fig. 4,5 | Fig. 4,5 | 80(2P) |  | F, L2/3 |  |  |
| PV+ ↔Pyr | not tested | not tested | not tested |  |  |  |  |
| Holmgren et al. 2003<br>(paired) |  |  |  | rat | V1, S, L2/3 | P14-16 | <100 μm<br>horizontal<br>offset |
| Pyr → PV+ | 243 | 121-182 | 50-75 |  |  |  |  |
| PV+ → Pyr | 243 | 121-170 | 50-70 |  |  |  |  |
| PV+ ↔Pyr | 243 | 60-126 | 25-52 |  |  |  |  |
| Kapfer et al. 2007<br>(multi-paired) |  |  |  | rat | S, L2/3 | P23±5<br>(AVG±STDEV) | <50 μm |
| Pyr → PV+ | 40 | 19 | 47.5 |  |  |  |  |
| PV+ → Pyr | 39 | 26 | 66.7 |  |  |  |  |
| PV+ ↔Pyr |  |  | Not reported |  |  |  |  |
| Campagnola et al. 2021<br>(multi-paired) |  |  |  | mouse | V1,L2/3-L6 | P46±4.6<br>(AVG±STDEV) | <100 μm |
| Pyr → PV+ | 50 | 21 | 42(L2/3) | | | | |
| PV+ → Pyr | 17 | 49 | 34.69(L2/3) | | | | |
| PV+ ↔ Pyr | Pooled inhibitory and normalized | | | | | | |
| Espinoza et al. 2018<br>(multi-paired) | | | | mouse | Dentate Gyrus, GC cells | P20-44 | <100 μm |
| GC → PV+ | not specified | not specified | 11 | | | | |
| PV+ → GC | 1301 | 296 | 22.75 | | | | |
| PV+ ↔ GC | 1301 | 32 | 2.46 | | | | |
| Frandolig et al. 2019<br>(paired) | | | | mouse | S1, L6a | P13-47 | <100 μm |
| (CThN)Pyr → PV+ | 86 | 32 | 37 | | Upper L6a | | |
| PV+ → Pyr (CThN) | 86 | 46 | 54 | | Upper L6a | | |
| PV+ ↔ Pyr (CThN) | 86 | 23 | 27 | | Upper L6a | | |
| (CCN)Pyr → PV+ | 78 | 34 | 44 | | Upper L6a | | |
| PV+ → Pyr(CCN) | 77 | 31 | 40 | | Upper L6a | | |
| PV+ ↔Pyr(CCN) | 77 | 21 | 27 |  | Upper L6a |  |  |
| (CThN)Pyr → PV+ | 46 | 9 | 20 |  | Lower L6a |  |  |
| PV+ → Pyr (CThN) | 46 | 17 | 37 |  | Lower L6a |  |  |
| PV+ ↔Pyr (CThN) | 46 | 6 | 13 |  | Lower L6a |  |  |
| (CCN)Pyr → PV+ | 33 | 14 | 42 |  | Lower L6a |  |  |
| PV+ → Pyr(CCN) | 33 | 14 | 42 |  | Lower L6a |  |  |
| PV+ ↔Pyr(CCN) | 33 | 7 | 21 |  | Lower L6a |  |  |
| Gabernet et al. 2005<br>(paired) |  |  |  | mouse | S1,L4 | P14-25 | Not reported |
| Pyr → PV+ | not<br>reported | not<br>reported | not<br>reported |  |  |  |  |
| PV+ → Pyr | not<br>reported | 6 | ~50% |  |  |  |  |
| PV+ ↔Pyr | not<br>reported | not<br>reported | not<br>reported |  |  |  |  |
| Gainey et al. 2018<br>(paired) |  |  |  | mouse | S1, L2/3 | P18-21 | <60 μm |
| Pyr → PV+ | not<br>tested | not tested | not<br>tested |  |  |  |  |
| PV+ → Pyr | 12 | 11 | 91.67 |  |  |  |  |
| PV+ ↔ Pyr | not tested | not tested | not tested |  |  |  |  |
| Gibbson et al 1999<br>(paired) |  |  |  | rat | S1, L4, L6 | P14-21 | <50 µm |
| RS → FS & LTS | 54 | 22 | 40.74 |  |  |  |  |
| FS & LTS → RS | 71 | 37 | 52.11 |  |  |  |  |
| FS & LTS ↔ RS | 54 | 5 | 9.26 |  |  |  |  |
| Guan et al. 2017<br>(multi-paired) |  |  |  | mouse | V1, L2/3 | P12-18 | <100 µm |
| Pyr → PV+ | not tested | not tested | not tested |  |  |  |  |
| PV+ → Pyr | 82<br>74 | 66<br>56 | 80.49<br>75.68 |  | (V1)<br>(Cg1/2) |  |  |
| PV+ ↔ Pyr | not tested | not tested | not tested |  |  |  |  |
| Gupta & Markram 2005<br>(multi-paired) |  |  |  | rat | S1, L2-4 | P13-16 | Not reported |
| Pyr → PV+ | not tested | not tested | not tested |  |  |  |  |
| FS → Pyr (depressing) | 131 | 100 | 76.3 |  |  |  |  |
| PV+ ↔ Pyr | not tested | not tested | not tested |  |  |  |  |
| Beierlein et al. 2002 |  |  |  | rat | S,L6 | P14-21 | <50 µm |
| (paired) |  |  |  |  |  |  |  |
| Pyr → FS | 41 | 4 | 9.76 | | | | |
| PV+ → Pyr | not<br>reported | not<br>reported | not<br>reported | | | | |
| PV+ ↔ Pyr | not<br>reported | not<br>reported | not<br>reported | | | | |
| Avermann et al. 2012<br>(multi-paired) | | | | mouse | S1, L2/3 | P17-22 | <160 μm |
| Pyr → PV+(FS) | Fig. 7 | Fig. 7 | 58 | | | | |
| (FS)PV+ → Pyr | Fig. 7 | Fig. 7 | 60 | | | | |
| (FS)PV+ ↔ Pyr | 36 | 12 | 33.3 | | | | |
| Hofer et al. 2011<br>(multi-paired) | | | | mouse | V1, L2/3 | adult, not<br>specified | < 50 μm |
| Pyr → PV+ | 41 | 36 | 87.81 | | | | |
| PV+ → Pyr | not<br>tested | not tested | not<br>tested | | | | |
| PV+ ↔ Pyr | not<br>tested | not tested | not<br>tested | | | | |
| House et al. 2011<br>(paired) | | | | rat | S1,L2/3 | P18-24 | < 150 μm |
| Pyr → FS | not<br>tested | not tested | not<br>tested | | | | |
| FS → Pyr | 45 | 21 | 46.7 | | | | |
| FS ↔ Pyr | not tested | not tested | not tested | | | | |
| Ali et al. 1999<br>(paired, sharp) | | | | rat | CA1 | Adults, not specified | < 200 μm |
| Pyr → FS (basket) | 263 | not reported | not reported | | | | |
| FS (basket) → Pyr | 263 | 55 | 20.91 | | | | |
| FS (basket) ↔ Pyr | 263 | 2 | 0.76 | | | | |
| Jouhannneau et al.<br>2018<br>(*in-vivo* multi-paired) | | | | mouse | S1, L2/3 | P21-30 | < 160 μm |
| Pyr → PV+ | 39 | 17 | 43.6 | | | | |
| PV+ → Pyr | 33 | 20 | 60.6 | | | | |
| PV+ ↔ Pyr | not reported | not reported | not reported | | | | |

